# Two bifunctional inositol pyrophosphate kinases/phosphatases control plant phosphate homeostasis

**DOI:** 10.1101/467076

**Authors:** Jinsheng Zhu, Kelvin Lau, Robert K. Harmel, Robert Puschmann, Larissa Broger, Amit K. Dutta, Henning J. Jessen, Ludwig A. Hothorn, Dorothea Fiedler, Michael Hothorn

**Affiliations:** Structural Plant Biology Laboratory, Department of Botany and Plant Biology, University of Geneva, Switzerland.; Leibniz-Forschungsinstitut für Molekulare Pharmakologie, Berlin, Germany & Department of Chemistry, Humboldt Universität zu Berlin, Berlin, Germany; Institute of Organic Chemistry, Albert-Ludwigs-Universität Freiburg, Germany; Institute of Biostatistics, Leibniz University, Hannover, Germany

## Abstract

Many eukaryotic proteins regulating phosphate (Pi) homeostasis contain SPX domains. We have previously shown that these domains act as cellular receptors for inositol pyrophosphate (PP-InsP) signaling molecules, suggesting that PP-InsPs may regulate Pi homeostasis. Here we report that simultaneous deletion of two diphosphoinositol pentakisphosphate kinases VIH1 and 2 in Arabidopsis impairs plant growth and leads to constitutive Pi starvation responses. We demonstrate that VIH1 and VIH2 are bifunctional cytosolic enzymes able to generate and break-down PP-InsPs. Point-mutants targeting the kinase and phosphatase active sites have opposing effects on plant Pi content and Pi starvation responses, while VIH1 and VIH2 protein levels remain constant in different Pi growth conditions. Enzymatic assays reveal that ATP-Mg^2+^ substrate levels can shift the relative kinase and phosphatase activities of full-length diphosphoinositol pentakisphosphate kinases. Deletion of phosphate starvation response transcription factors rescues *vih1 vih2* mutant phenotypes, placing diphosphoinositol pentakisphosphate kinases and PP-InsPs in plant phosphate signal transduction cascades. We propose that VIH1 and VIH2 relay changes in cellular ATP concentration to changes in PP-InsP levels, allowing plants to maintain cellular Pi concentrations constant and to trigger Pi starvation responses.

## Introduction

Phosphorus is a growth limiting nutrient for plants, taken up from the soil as inorganic phosphate (H_2_PO_4_^-^, Pi). Plants and other soil-living eukaryotes have evolved sophisticated Pi sensing, uptake, transport and storage mechanisms. Many of the proteins involved in these processes contain small hydrophilic SPX domains (Secco et al., 2012). We have previously shown that these domains act as cellular receptors for inositol pyrophosphate (PP-InsP) signaling molecules (Wild et al., 2016). PPInsPs are composed of a fully phosphorylated inositol ring containing one or two pyrophosphate moieties (Shears, 2018). PP-InsPs bind to a basic surface cluster highly conserved among SPX domains to regulate divers biochemical and cellular processes in fungi, plants and animals. The N-terminal SPX domains of the yeast VTC complex stimulates its inorganic polyphosphate polymerase activity when bound to 5PP-InsP_5_ (Hothorn et al., 2009; Wild et al., 2016; Gerasimaite et al., 2017). In plant and human phosphate exporters, mutations in the PP-InsP binding surfaces of their N-terminal SPX domains affect their Pi transport activities (Legati et al., 2015; Wild et al., 2016). Plants contain additional soluble, stand-alone SPX proteins, which have been shown to bind PHOSPHATE STARVATION RESPONSE (PHR) transcription factors (Rubio et al., 2001; Puga et al., 2014; Wang et al., 2014; Qi et al., 2017). PHR1 and its homolog PHL1 are master regulators of the plant Pi starvation response, which enables plants to grow and survive in Pi limiting growth conditions (Bustos et al., 2010). PP-InsP-bound SPX domains can physically interact with PHR1, keeping it from binding DNA (Puga et al., 2014; Wang et al., 2014; Wild et al., 2016; Qi et al., 2017). In the absence of PP-InsPs, the SPX – PHR1 complex dissociates and PHR1 oligomers can target the promoters of Pi starvation induced (PSI) genes, resulting in major changes in plant metabolism and development (Wild et al., 2016; Jung et al., 2018). These findings suggest that changes in cellular PP-InsP levels may regulate eukaryotic Pi homeostasis. 5PP-InsP_5_ levels are indeed reduced in yeast and in animal cells grown in Pi starvation conditions (Lonetti et al., 2011; Wild et al., 2016; Gu et al., 2017a), suggesting that PP-InsP metabolic enzymes may be key regulators of Pi homeostasis (Gu et al., 2017a; Azevedo and Saiardi, 2017; Shears, 2018).

PP-InsP biosynthesis is well understood in yeast and mammals, where inositol hexakisphosphate kinases Kcs1/IP6K catalyze the synthesis of 5PP-InsP_5_ from InsP_6_. 5PP-InsP_5_ can be further phosphorylated by diphosphoinositol pentakisphosphate kinases Vip1/PPIP5K to 1,5(PP)_2_-InsP_4_ (referred as InsP_8_ in this study) (Shears, 2018) (Figure 1). How these enzymes may sense changes in cellular Pi concentration remains to be elucidated in mechanistic detail. Currently it is known that Kcs1/IP6Ks have a K_m_ for ATP in the low millimolar range and may therefore generate 5PP-InsP5 in response to changes in cellular ATP levels (Saiardi et al., 1999; Gu et al., 2017a, 2017b). Vip1/PPIP5Ks are bifunctional enzymes harboring an N-terminal InsP kinase and and a C-terminal phosphatase domain (Figure 2A) (Mulugu et al., 2007; Fridy et al., 2007; Wang et al., 2015). The yeast and animal kinase domain acts on the phosphate group at the C1 position of the inositol ring of both InsP_6_ and 5PP-InsP_5_, yielding 1PP-InsP_5_ or InsP_8_ reaction products (Mulugu et al., 2007; Fridy et al., 2007). The phosphatase domain has been characterized as a specific 1PP-InsP_5_ / InsP_8_ phosphatase in yeast (Wang et al., 2015). Deletion of PPIP5K in human cells results in lower InsP_8_ and elevated ATP levels, due to enhanced mitochondrial oxidative phosphorylation and increased glycolysis (Gu et al., 2017b).

**Figure 1.**
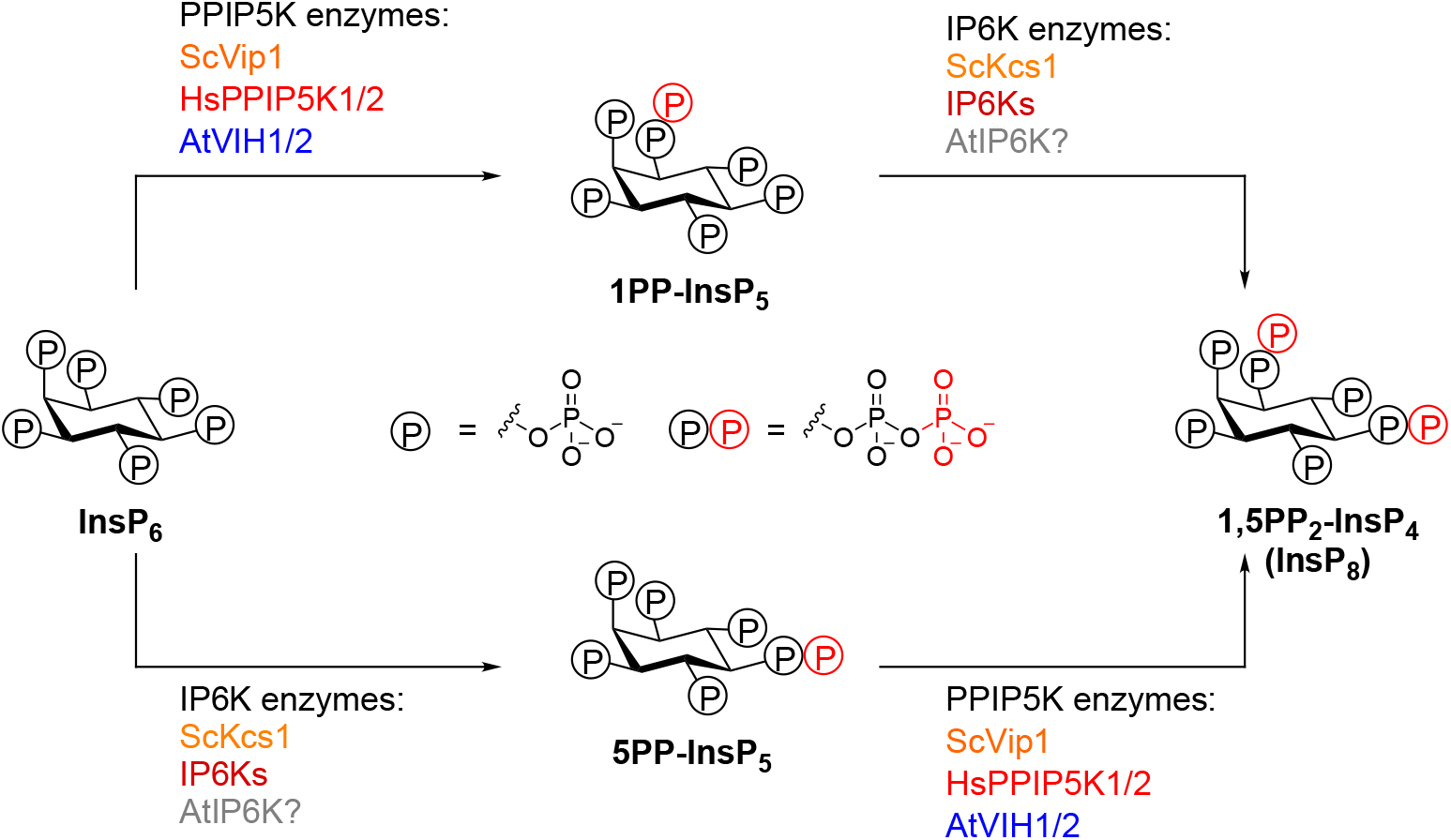
Overview of PP-InsP isoforms and kinases involved in inositol pyrophosphate metabolism. Two classes of inositol pyrophosphate kinases, PPIP5K and IP6K, are responsible for the synthesis of inositol pyrophosphates. The schematic describes PPIP5K enzymes from *Saccharomyces cerevisiae* ScVip1, *Homo sapiens* HsPPIP5K11 and 2, *Arabidopsis thaliana* VIH1 and VIH2. These enzymes synthesize 1PP-InsP_5_ and 1,5(PP)_2_-InsP_4_ (InsP_8_) by phosphorylating InsP_6_ at the 1 position, and by phosphorylating 5PP-InsP_5_ at the 1 position, respectively. IP6K enzymes from *Saccharomyces cerevisiae* KCS1 and *Homo sapiens* IP6Ks synthesize 5PP-InsP_5_ and 1, 5(PP)_2-_InsP_4_ (InsP_8_) by phosphorylating InsP_6_ at the 5 position, and by phosphorylating 1PP-InsP_5_ at the 5 position, respectively. However, plant IP6Ks have not been reported yet.

In plants no IP6K enzyme has been identified thus far, but Vip1/PPIP5K orthologs Vip1 and VIH1/2 have been reported from Chlamydomonas (Couso et al., 2016) and Arabidopis (Desai et al., 2014; Laha et al., 2015), respectively. The algal and plant enzymes share the bifunctional kinase/phosphatase domain architecture with yeast and animal PPIP5Ks and consequently Arabidopsis VIH1 and VIH2 can rescue yeast *vip1* but not *ksc1* mutants (Desai et al., 2014; Laha et al., 2015). Deletion of the Chlamydomonas Vip1 results in nutrient signaling phenotypes affecting carbon metabolism (Couso et al., 2016). The Arabidopsis *vih2-4* loss-of-function mutant has lower InsP_8_ but increased InsP_7_ levels, suggesting that the enzyme may generate a InsP_8_ isoform *in vivo* (Laha et al., 2015). VIH2 mutant lines have been reported to affect jasmonic acid-regulated herbivore resistance, as PP-InsPs also form critical co-factors of the jasmonate receptor complex (Sheard et al., 2010; Laha et al., 2016). No obvious Pi starvation phenotype has been reported for these mutants (Kuo et al., 2018). To analyze if indeed plant Pi homeostasis is mediated by PP-InsPs, we here characterize Pi signaling related phenotypes of *vih1* and *vih2* mutants in Arabidopsis and dissect the individual contributions of their conserved InsP kinase and phosphatase domains to Pi homeostasis and Pi starvation responses.

## Results

### Deletion of VIH1/VIH2 affects plant growth and phosphate homeostasis

Arabidopsis VIH1 and VIH2 show 89 % sequence identity at the protein level (Figure 2 – figure supplement 2). VIH1 has been previously reported to be specifically expressed in pollen while VIH2 showed a broad expression pattern as judged from qRT-PCR experiments (Desai et al., 2014; Laha et al., 2015). We further analyzed *VIH1* and *VIH2* expression in Arabidopsis using pVIH1::GUS and pVIH2::GUS reporter lines. We found *VIH2* to be broadly expressed in different tissues and organs, and strong expression for *VIH1* in pollen as previously reported (Laha et al., 2016) (Figure 2 – figure supplement 2). However, our GUS lines revealed additional expression of *VIH1* in root tips, where also *VIH2* is expressed (Figure 2 – figure supplement 2). VIH1 and VIH2 reporter lines harboring a C-terminal mCitrine (mCit) tag (which complements the *vih1 vih2* mutant phenotype, compare Figure 3A, see below) revealed that VIH1 is present in pollen grains, pollen tubes and in the root tip, where also VIH2 is expressed (Figure 2 – figure supplement 2). Thus VIH1 and VIH2 show partially overlapping expression in different tissues and organs and both enzymes are localized in the cytoplasm (Figure 2 – figure supplement 2; compare Figure 5E). This prompted us to generate *vih1 vih2* double mutants using the *vih1-2*, *vih1-6* and *vih2-4* T-DNA insertion lines as knock-out mutants of the respective enzyme (Figure 2A, B).

**Figure 2.**
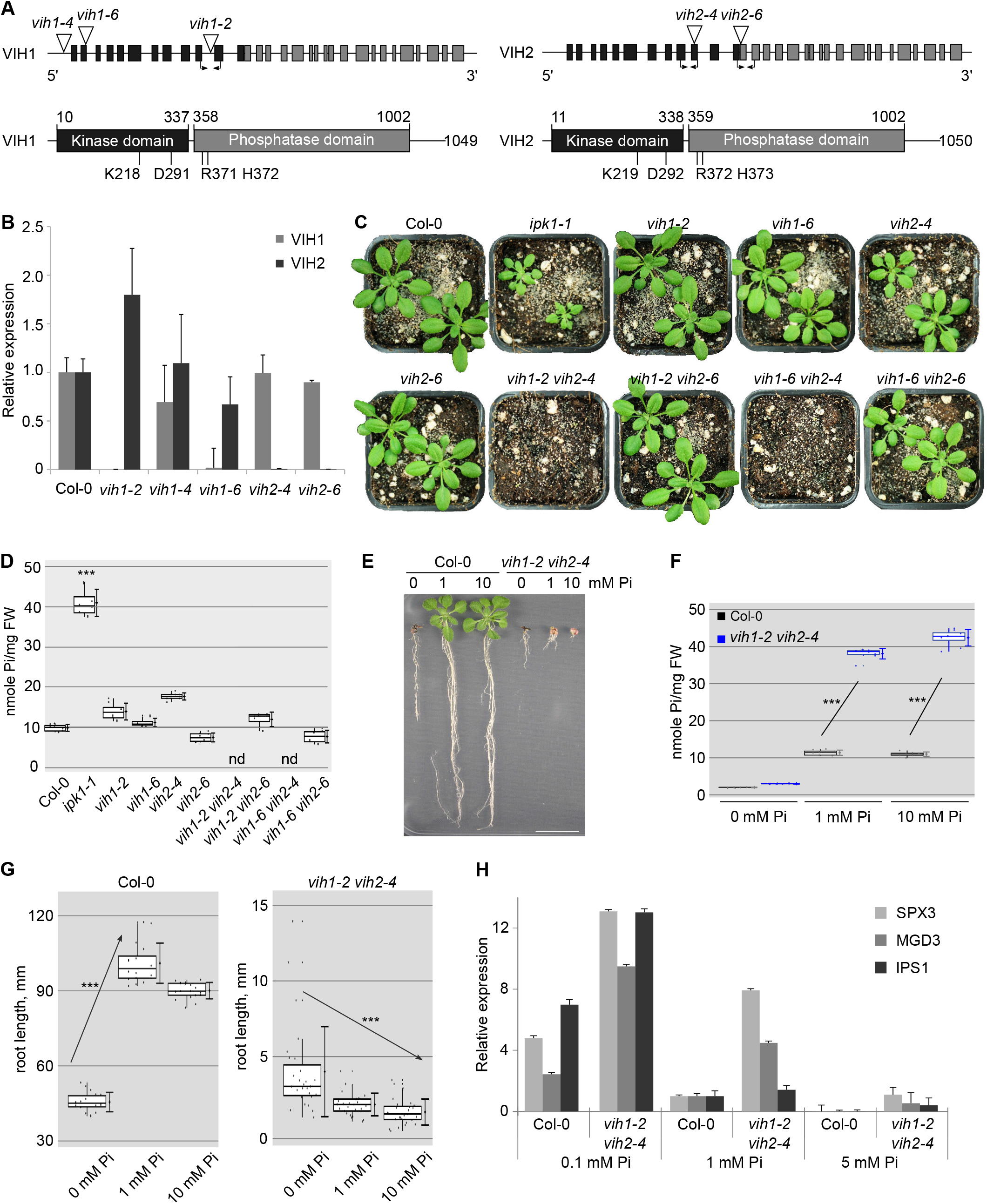
*vih1 vih2* loss-of-function mutants show severe growth phenotypes and hyperaccumulate phosphate. **(A)** Schematic overview of VIH1 and VIH2: (upper panel) VIH1 and VIH2 genes with exons described as rectangles, introns as lines. T-DNA insertions are depicted as triangles, primer positions used in qRT-PCR analyses are indicated by arrows. (lower panel) VIH1 and VIH2 protein architecture, with kinase domains shown in black, phosphatase domains in gray and connecting linkers and putatively unstructured regions as lines. The positions of the catalytic point mutations used in this study are shown alongside. **(B)** qRT-PCR expression analysis of VIH1 and VIH2 transcripts in the T-DNA mutant allele backgrounds. Averages of triplicate reactions ± SE are shown. Quantifications were repeated at least 3 times independently with similar results. **(C)** Growth phenotypes of *ipk1*, *vih1* and *vih2* single mutant, and of *vih1 vih2* double mutants 20 DAG in soil compared to Col-0 wild-type. *ipk1-1* represents a mild loss-of-function mutant allele of Arabidopsis inositol pentakisphosphate 2-kinase. **(D)** Shoot Pi contents of the plants shown in (B). For each boxed position, 4 independent plants were measured with 2 technical replicates. **(E)** Root growth phenotypes of Col-0 wild-type and *vih1-2 vih2-4* seedlings grown in different Pi concentrations. Plants were germinated in vertical ^1/2^MS plates for 8 d, transferred into Pi-deficient^1/2^MS plates supplemented with either 0 mM, 1 mM or 10 mM Pi and grown for additional 12 d.Scale bars correspond to 2 cm. (**F**) Shoot Pi content of the wild-type (black) and *vih1-2 vih2-4* (blue) seedlings grown under different Pi conditions as shown in (E). For each boxed position, at least 3 independent measurements were performed for shoots of seedlings from 3 different MS plates. (**G**) Trend analysis of seedling root length vs. phosphate concentration for the seedlings described in(E) Col-0 wild-type roots are shown on the left side, *vih1-2 vih2-4* roots are plotted on right side. For each boxed position, at least 16 independent measurements were performed for roots of seedlings from 3 different MS plates. **(H)** qRT-PCR quantification of three phosphate starvation induced (PSI) marker genes SPX3, MGD3 and IPS1, in wild-type and *vih1-2 vih2-4* seedlings 12 DAG grown in plates supplemented with 0.1 mM, 1 mM and 5 mM Pi for 5 days.

We found that *vih1-2 vih2-4* and *vih1-6 vih2-4* double mutants displayed severe growth phenotypes, while *vih1* and *vih2* single mutants and *vih1-2 vih2-6* or *vih1-6 vih2-6* double mutants looked similar to wild-type plants (Figure 2C). In line with this, *vih2-4* plants expressing an inducible artificial microRNA (amiRNA) targeting the VIH1 transcript showed intermediate growth phenotypes (Figure 2 – figure supplement 2). Taken together, independent genetic experiments suggest that VIH1 and VIH2 act redundantly in Arabidopsis PP-InsP metabolism, and that simultaneous loss-of-function of these conserved enzymes strongly affects plant growth.

We next studied phosphate-related phenotypes in our mutant backgrounds. We found shoot Pi levels to be significantly increased in *ipk1* (Kuo et al., 2014) but not in *vih1* or *vih2* single mutants or *vih1-2 vih2-6* or *vih1-6 vih2-6* double mutants, when compared to wild-type (Figure 2C). Due to their strong growth phenotype, we could not assess Pi shoot contents for adult *vih1-2 vih2-4* or *vih1-6 vih2-4* double mutants (Figure 2C, D). We thus grew *vih1-2 vih2-4* heterozygous mutants on half-strength MS (^1/2^MS) plates and transferred seedlings exhibiting the double mutant phenotype to plates supplemented with different concentrations of Pi (Figure 2E). We found that *vih1-2 vih2-4* double mutant seedlings accumulated significantly more Pi than wild-type plants, when grown in 1 mM Pi and 10 mM Pi conditions, but not in 0 mM Pi condition (Figure 2F). While wild-type seedlings showed a decrease in root length in response to Pi limitation, *vih1-2 vih2-4* mutant seedlings showed the opposite trend (Figure 2G). We hypothesized that this phenotype may be due to *vih1-2 vih2-4* plants lacking a PP-InsP isoform critically involved in the Pi starvation response (Wild et al., 2016; Jung et al., 2018). Consistently, we found PSI gene expression to be up-regulated in the double mutant, possibly indicating that PHR1 / PHL1 may not be under the negative regulation by PP-InsP-bound SPX domains (Wild et al., 2016; Qi et al., 2017) (Figure 2H). In line with this, estradiol-induced expression of an amiRNA targeting the *VIH1* transcript in the *vih2-4* single mutant background leads to a significant increase in shoot Pi content when compared to the uninduced control (Figure 2 – figure supplement 2). Together, these experiments suggest a role for PP-InsPs in Pi homeostasis and starvation responses.

### VIH1 and 2 are part of the plant phosphate starvation response pathway

We next complemented *vih1-2 vih2-4* mutant lines with VIH1/VIH2 wild-type constructs containing a C-terminal mCit(rine) fluorescent tag (Figure 3A; Figure 2 – figure supplement 2). Expression of wild-type VIH2 from its endogenous promoter fully rescued the double mutant growth phenotype and shoot Pi content (Figure 3A-C). For VIH1, we observed lines showing partial complementation with reduced protein levels, increased shoot Pi concentrations, and induction of PSI genes (Figure 3A-D). We could not complement the *vih1-2 vih2-4* mutant phenotype with constructs expressing a VIH2^KD/AA^ version containing point-mutations targeting the kinase domain (Figure 3E; Figure 3 – figure supplement 3B). In contrast, expression of a construct harboring point-mutations in the phosphatase domain (VIH2^RH/AA^) at least partially recovers the *vih1-2 vih2-4* development and growth phenotypes (Figure 3E,F; Figure 3 – figure supplement 3). VIH2^RH/AA^ lines show significantly reduced shoot Pi levels but no drastic changes in PSI gene expression (Figure 3G,H). We next analyzed the *vih2-6* allele, also recently described as *vip1-1* (Kuo et al., 2018). We found that *vih2-6* results in a shortened transcript, which may encode a truncated version of VIH2 harboring the PPIK5K kinase domain only (Figure 3 – figure supplement 3). Notably, *vih2-6* mutant plants show significantly reduced shoot Pi levels and reduced PSI gene expression, when compared to wild-type (Figure 3I-L). These specific phenotypes can be rescued by expressing wild-type VIH2 in the *vih2-6* mutant background (Figure 3I-L). Taken together, our complementation analyses reveal an essential role for the VIH kinase domain in plant growth and Pi signaling, while the VIH phosphatase domain may have a regulatory function in plant Pi homeostasis and signaling.

**Figure 3.**
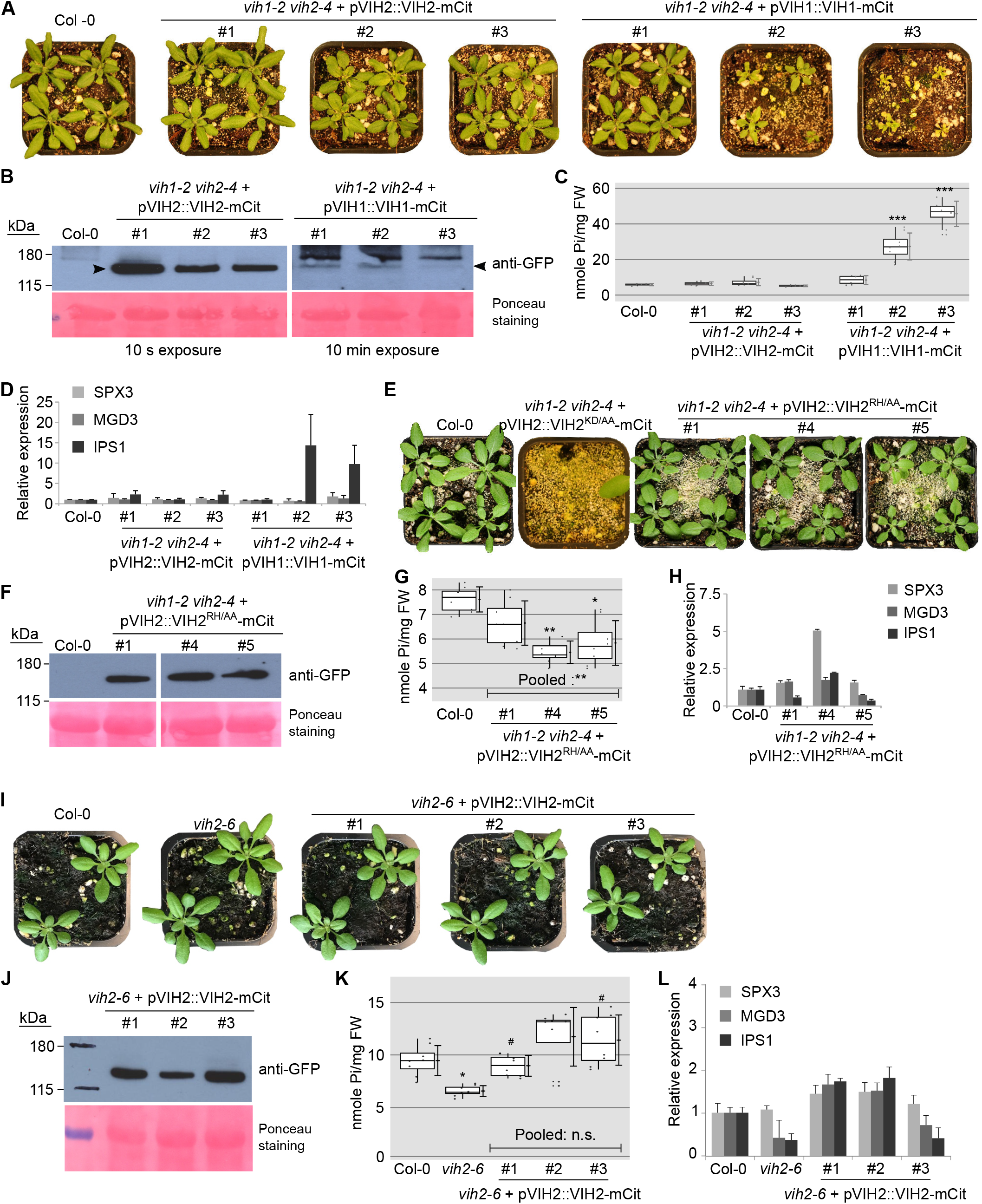
VIH kinase and phosphatase activities regulate plant Pi homeostasis. **(A)** Expression of VIH1-mCit or VIH2-mCit under the control of their endogenous promoter can complement the *vih1-2 vih2-4* phenotype. Seedlings 7 DAG were transferred to soil and grown for 23 d. **(B)** Western blot showing the expression of VIH1-mCit and VIH2-mCit proteins (indicated by arrow heads) in the transgenic lines from (A) using an anti-GFP antibody. A Ponceau stain of the membrane is shown as loading control below. **(C)** Shoot Pi content of plants 20 DAG. For each position, 4 independent plants were measured with 2 technical replicates each. **(D)** qRT-PCR quantification of the PSI marker genes SPX3, MGD3 and IPS1, in Col-0 wild-type and *vih1-2 vih2-4* seedlings 10 DAG. **(E)** Expression of pVIH2::VIH2^RH/AA^-mCit (Arg372Ala and His373Ala) but not pVIH2::VIH2^KD/AA-^mCit (Lys219Ala and Asp298Ala) rescues the *vih1-2 vih2-4* mutant phenotype. Plants were transferred to soil 7 DAG and grown for 20 d. **(F)** Western blot showing the VIH2^RH/AA^-mCit protein levels in the complementation lines described in (E), detected using an anti-GFP antibody. **(G)** Shoot Pi contents of plants 20 DAG described in (E). For each boxed position, 4 independent plants were measured with 2 technical replicates each. **(H)** qRT-PCR quantification of the PSI marker genes SPX3, MGD3 and IPS1, in wild-type and *vih1-2 vih2-4* seedlings 10 DAG complemented with pVIH2::VIH2^RH/AA^-mCit. **(I)** Growth phenotypes of the *vih2-6* mutant expressing the kinase domain-only, and 3 independent lines of *vih2-6* complemented with pVIH2::VIH2-mCit. Plants 7 DAG were transferred to soil and grown for 21 d. **(J)** Western blot of the complementation lines shown in (I). The VIH2-mCit fusion protein was detected using an anti-GFP antibody. **(K)** Shoot Pi content of plants 20 DAG as described in (I). For each position, 4 independent plants were measured with 2 technical replicates each. **(L)** qRT-PCR quantification of the PSI marker genes SPX3, MGD3 and IPS1, in wild-type and *vih2-6* seedlings 10 DAG complemented with pVIH2::VIH2-mCit.

To test our hypothesis that diphosphoinositol pentakisphosphate kinases are part of the plant Pi starvation response, we next analyzed the genetic interaction between VIH1, VIH2 and the known phosphate starvation response transcription factors PHR1 and PHL1 (Bustos et al., 2010). We found at seedling stage that a *vih1-2 vih2-4 phr1* triple mutant partially restores the seedling growth phenotype of the *vih1-2 vih2-4*, but still hyper-accumulate phosphate under Pi sufficient growth conditions, similar to the *vih1-2 vih2-4* double mutant (Figure 4A,B). The *vih1-2 vih2-4 phr1 phl1* quadruple mutant fully restores root growth and partially complements the shoot phenotypes with shoot and root Pi levels being similar to wild-type (Figure 4A,B). Importantly the *vih1-2 vih2-4* root growth phenotype under different Pi regimes (compare Figure 2G, see above) is restored in the triple and quadruple mutants (Figure 4C).

**Figure 4.**
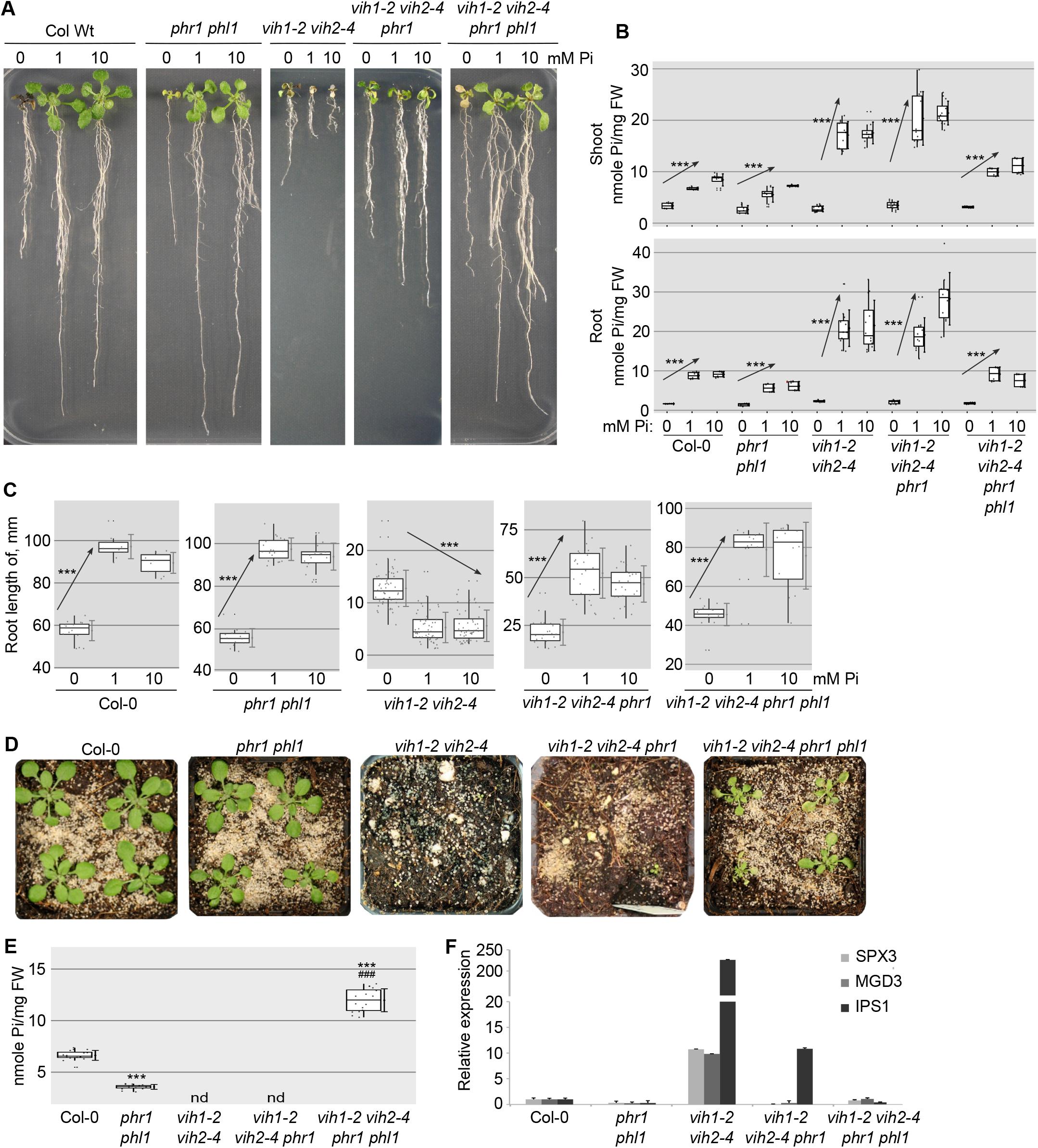
Mutation of PHR1 and PHL1 rescues *vih1-2 vih2-4* mutant phenotypes. **(A)** Growth phenotype of the *phr1 phl1* double mutant, *vih1-2 vih2-4* double mutant, *vih1-2 vih2-4 phr1* triple mutant, and *vih1-2 vih2-4 phr1 phl1* quadruple mutant grown in different Piconcentrations. The triple and quadruple mutants were obtained by crossing *phr1 phl1* homozygous plants with *vih1-2 vih2-4* heterozygous plants. Seedlings 7 DAG were transplanted from ^1/2^MS plates to Pi-deficient ^1/2^MS liquid supplemented with 0 mM, 1 mM or 10 mM Pi, and grown for additional 7 d. **(B)** Shoot Pi contents (top panel) and root Pi contents (bottom panel) of seedlings 14 DAG as described in (A). For each Pi concentration, at least 4 independent plants were measured with 2 technical replicates each. **(C)** Root length of seedlings 14 DAG as described in (A). For each Pi concentration, seedlings from at least 3 independent plates were measured. **(D)** Growth phenotype of *phr1 phl1* double mutant, *vih1-2 vih2-4* double mutant, *vih1-2 vih2-4 phr1* triple mutant, and *vih1-2 vih2-4 phr1 phl1* quadruple mutant. Seedlings 7 DAG were transferred to soil and grown for 21 d. **(E)** Shoot Pi content of plants 20 DAG as described in (D). 4 independent plants were measured with 2 technical replicates each. **(F)** qRT-PCR quantification of the expression of PSI genes, using mRNA extracted from seedlings 10 DAG.

At later developmental stages, only the *vih1-2 vih2-4 phr1 phl1* quadruple mutant could partially rescue the *vih1-2 vih2-4* phenotype and restore normal PSI gene expression (Figure 4D, F). Shoot phosphate levels in the quadruple mutant are significantly higher when compared to either wild-type or *phr1 phl1* double mutants (Figure 4E). This suggests that VIH1 and VIH2 generated PP-InsPs regulate the activity not only of PHR1 and PHL1, but possibly of other Pi starvation responsive transcription factors such as PHL3 and/or PHL4 (Sun et al., 2015; Wang et al., 2018) or of other PP-InsP responsive Pi signaling components (Hamburger et al., 2002; Wild et al., 2016). Together, our analysis firmly places VIH1 and VIH2 in the plant phosphate starvation response pathway.

### VIH kinase and phosphatase domains together control plant Pi homeostasis

We speculate that concerted VIH2 kinase and phosphatase activities may enable plants to maintain Pi homeostasis by controlling the cellular levels of PP-InsPs. In line with this, plants constitutively over-expressing VIH2 in wild-type background resemble wild-type plants, while over-expression of either VIH2^KD/AA^ or VIH2^RH/AA^ results in severe growth defects (Figure 5A,B, Figure 5 – figure supplement 5). Shoot Pi levels in VIH2^KD/AA^ are significantly higher compared to wild-type, while VIH2^RH/AA^ plants show lower Pi levels (Figure 5C). This suggest that the kinase and phosphatase activities of VIHs regulate Pi homeostasis in an antagonistic fashion. It is of note however that PSI target gene expression is misregulated in both VIH2^KD/AA^ and VIH2^RH/AA^ over-expression lines (Figure 5D).

**Figure 5.**
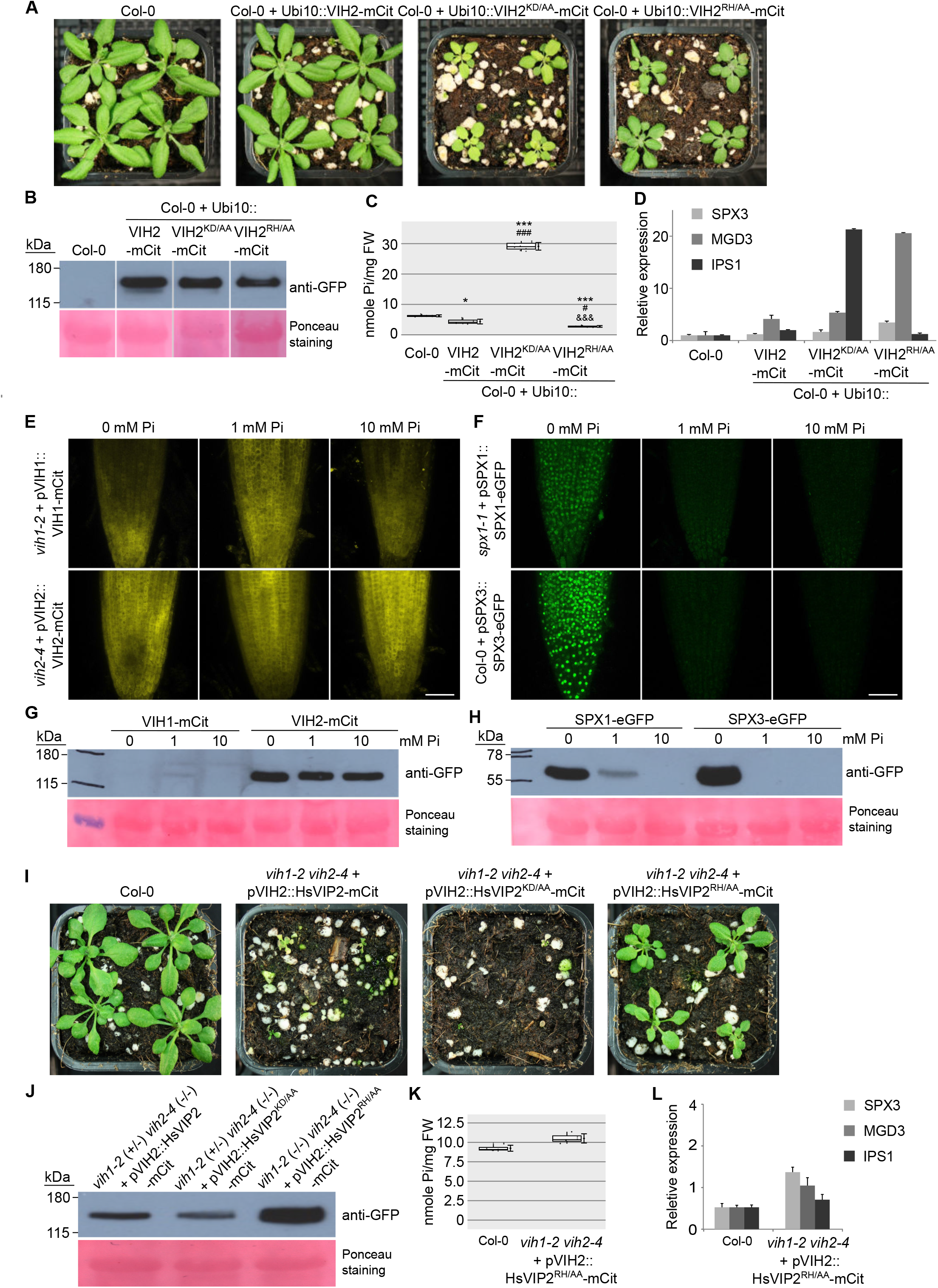
The kinase and phosphatase activities of VIH1 and VIH2 together control plant Pi homeostatis. **(A)** Growth phenotypes of plants over-expressing VIH2-mCit, VIH2^KD/AA^-mCit or VIH2^RH/AA^-mCit with a Ubi10 promoter in the Col-0 wild-type background. Seedlings 7 DAG were transferred to soil and grown for 21 d. **(B)** The expression of VIH2-mCit, VIH2^KD/AA^-mCit and VIH2^RH/AA^-mCit protein in plants as shown in (A) as detected by an anti-GFP antibody. **(C)** Shoot Pi content of plants 20 DAG as described in (A). 4 independent plants were measured with 2 technical replicates each. **(D)** qRT-PCR quantification of the expression of PSI genes in seedlings 10 DAG of Col-0 wild-type and VIH2-mCit, VIH2^KD/AA^-mCit and VIH2^RH/AA^-mCit over-expressed lines. **(E)** Representative confocal scanning microscopy images showing a mCit fluoresecent signal in the root tip of *vih1-2* and *vih2-4* seedlings transformed with pVIH1::VIH1-mCit or pVIH2::VIH2-mCit respectively. Seedlings 7 DAG were transferred from ^1/2^MS plates to Pi-deficient ^1/2^MS liquid medium supplemented with 0 mM, 1 mM or 10 mM Pi, and grown for 3 d. Scale bar = 50 μm. **(F)** Representative confocal scanning microscopy images showing a GFP fluorescent signal in the root tip of *spx1-1* and Col-0 wild-type seedlings transformed with pSPX1::SPX1-eGFP and pSPX3::SPX3-eGFP respectively. Seedlings were treated as described in (E). Scale bar = 50 μm. **(G)** Protein levels of VIH1-mCit and VIH2-mCit in plants grown in Pi-deficient media supplemented with 0 mM, 1 mM or 10 mM Pi and detected by an anti-GFP antibody. **(H)** Protein levels of SPX1-eGFP and SPX3-eGFP in plants grown grown in Pi-deficient media supplemented with 0 mM, 1 mM or 10 mM Pi and detected by an anti-GFP antibody. **(I)** Growth phenotype of *vih1-2 vih2-4* double mutants complemented with HsVIP2-mCit, HsVIP2^KD/AA^-mCit and HsVIP2^RH/AA^-Cit under the control of the AtVIH2 promoter. Seedlings 7 DAG are transferred to soil and grown for 21 d. **(J)** Expression of HsVIP2-mCit, HsVIP2^KD/AA^-mCit and HsVIP2^RH/AA^-Cit proteins, detected by an anti-GFP antibody. **(K)** Shoot Pi content of plants 20 DAG as described in (I). 4 independent plants were measured with 2 technical replicates each. **(L)** qRT-PCR quantification of the expression of PSI genes in seedlings 10 DAG of Col-0 wild-type and *vih1-2 vih2-4* mutant lines complemented with HsVIP2, HsVIP2^KD/AA^ and HsVIP2^RH/AA^.

We next tested if VIH1 and 2 promoter activities and protein levels change in response to different Pi growth regimes. We generated pVIH2::H(istone)2b-mCit lines, which accumulate the fluorescence signal in the nucleus, in either wild-type or *vih2-4* mutant background and found no significant changes in promoter activity in low to high Pi growth condition (Figure 5 – figure supplement 5A). In contrast, the known PSI targets SPX1 and SPX3 are highly induced under Pi starvation (Figure 5 – figure supplement 5A) (Duan et al., 2008). In agreement with these findings, our GUS reporter lines (compare Figure 2 – figure supplement 2) show slightly reduced, not increased VIH1 and VIH2 promoter activities under Pi starvation conditions (Figure 5 – figure supplement 5B). VIH1-mCit and VIH2-mCit fusion proteins expressed from their own promoter are expressed to similar levels in plants grown in different Pi conditions (Figure 5E,G), while the SPX1 and SPX3 proteins accumulate under Pi starvation (Figure 5F,H). Together, these experiments indicate that VIH enzyme activity may not be simply controlled at the transcript or protein level, but possibly at the level of the enzymatic activities of the kinase and phosphatase domains.

It has been previously suggested for human PPIP5K that the kinase and phosphatase domains may respond to changes in cellular ATP and Pi levels, respectively (Gu et al., 2017b; Shears, 2018). To test if the regulation of plant Pi homeostasis may be similar to the human system, we complemented our *vih1-2 vih2-4* mutant with the HsVIP2, codon-optimized for Arabidopsis and expressed under the control of the Arabidopsis VIH2 promoter (Figure 5I,J). We found that expression of wild-type HsVIP2 or HsVIP2^KD/AA^ carrying point-mutations that target the kinase domain, did not rescue the *vih1-2 vih2-4* mutant phenotype (Figure 5I,J; compare Figure 3E). Strikingly, expression of a HsVIP2^RH/AA^mutant in the phosphatase domain complemented the *vih1-2 vih2-4* mutant, resulting in wild-type-like shoot Pi levels and PSI target gene expression (Figure 5I-L). This indicates that human and plant PPIP5Ks may generate similar PP-InsP signaling molecules, able to regulate Pi homeostasis in these different organisms (Gu et al., 2017a). The absolute ATP, Pi and PP-InsP levels and PPIP5K kinase and phosphatase activities may however differ substantially between different organisms.

### Arabidopsis VIH2 is a 1-InsP kinase

The enzymatic properties of Arabidopsis VIH1 and VIH2 have not been characterized thus far. We mapped the catalytic PPIP5K kinase and phosphatase domains using the program HHPRED (Zimmermann et al., 2017) and expressed the isolated domains in insect cells (Figure 2 – figure supplement 2, see Methods). Next, we determined the substrate specificity of the VIH2 kinase domain (VIH2-KD, residues 11-338, Figure 7 – figure supplement 7A, C) in quantitative NMR assays using ^13^C-labeled InsP substrates (see Methods). In our assays we found that Arabidopsis VIH2-KD catalyzes the synthesis of 1PP-InsP_5_ from InsP_6_ and InsP_8_ from 5PP-InsP_5_ (Figure 6A,B; Figure 6 – figure supplement 6). We record similar activities for human PPIP5K2 (Figure 6A,B; Figure 7 – figure supplement 7A, C), in agreement with earlier reports (Choi et al., 2007; Fridy et al., 2007; Lin et al., 2009; Wang et al., 2011). The VIH2-KD^KD/AA^ mutant protein (Figure 7 – figure supplement 7A, C) shows no detectable activity using InsP_6_ as a substrate, and a minuscule activity towards 5PP-InsP_5_ (Figure 6C,D). NMR-derived enzyme kinetics suggest 5PP-InsP_5_ to be a better substrate for VIH2 when compared to InsP_6_ (Figure 6E,F). Together with the *in planta* observation that InsP_8_ levels are reduced and InsP_7_ levels are increased in *vih2-4* mutants (Laha et al., 2015), our data argue that the Arabidopsis VIH2 kinase domain may generate InsP_8_ *in vivo*.

**Figure 6.**
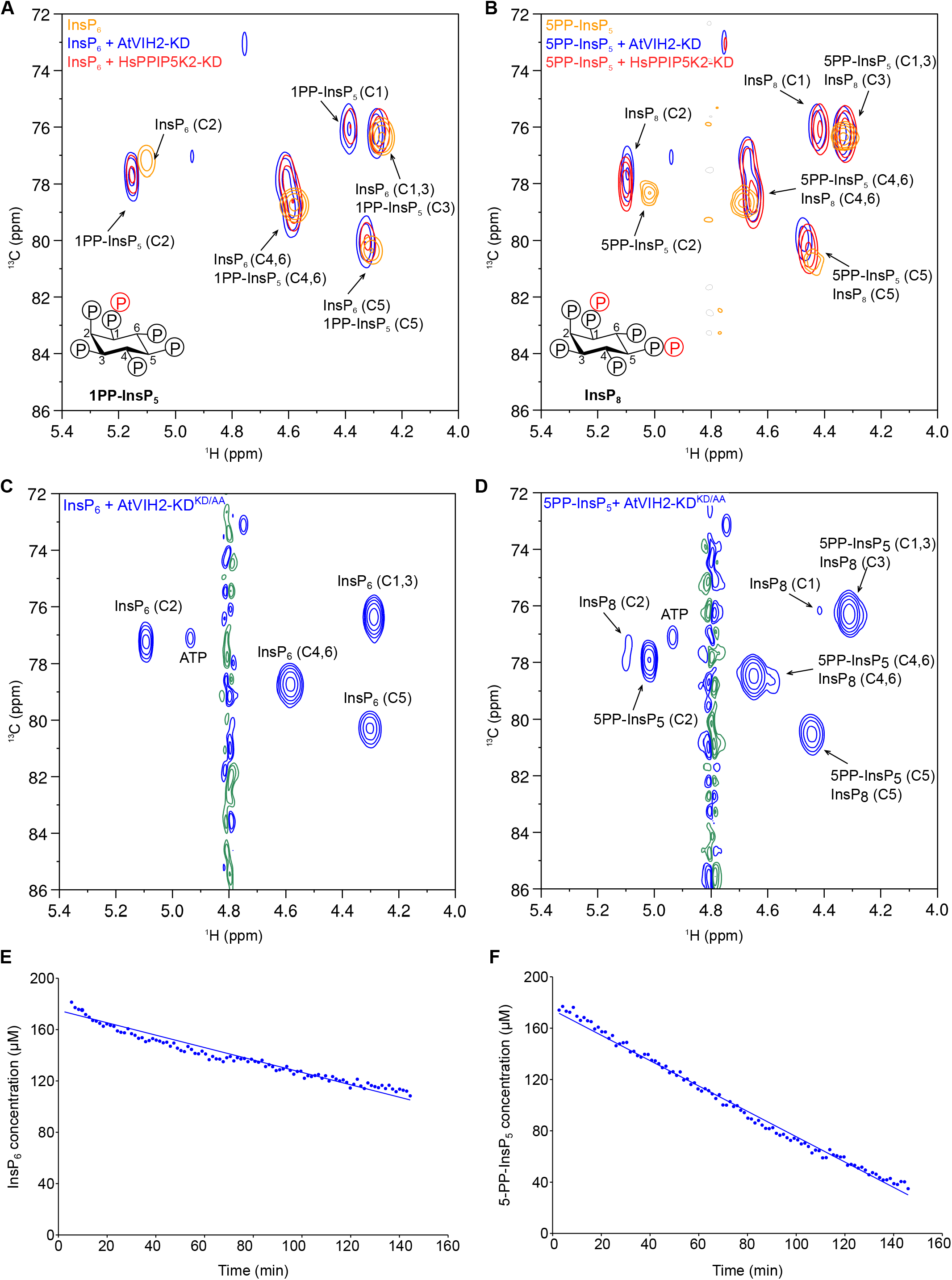
AtVIH2-KD displays 1-kinase activity and produces 1PP-InsP_5_ and InsP_8_. **(A, B)** 2D ^1^H-^13^C-HMBC spectra of the products produced by plant AtVIH2-KD (blue trace) and human HsVIP2-KD (red trace) in the presence of InsP_6_ (A) or 5PP-InsP_5_ (B). Substrate standards are colored in yellow. **(C, D)** 2D ^1^H-^13^C-HMBC spectra of the products produced by plant AtVIH2-KD_KD/AA_ (blue trace) in the presence of InsP_6_ (C) or 5PP-InsP_5_ (D). **(E, F)** Decay of the InsP_6_ (E) or 5-PP-InsP_5_ (F) substrate during a NMR-time course experiment. A fit of the initial decay indicates a turnover number of ~0.4/min with InsP_6_ as a substrate and ~1/min using 5-PP-InsP_5_ as a substrate.

### The VIH2 phosphatase domain has 1 and 5PP-InsPase activity

PPIP5Ks from different organisms all contain a C-terminal phosphatase domain (Pascual-Ortiz et al., 2018; Shears et al., 2017). Inositol pyrophosphatase activity has been established for fission yeast Asp1 and for human PPIP5K1 (Wang et al., 2015). The insect cell expressed and purified VIH2 phosphatase domain (VIH2-PD, residues 359-1002, Figure 7 – figure supplement 7B, C) hydrolyzes both 1PP-InsP_5_ and 5PP-InsP_5_, yielding InsP_6_, but not the non-hydrolyzable 5PCP-InsP_5_ analog (Figure 7A, B, C) (Wu et al., 2016). The *Saccharomyces cerevisiae* Vip1 phosphatase domain (Figure 7 – figure supplement 7B, C), which shares only 24% sequence identity with the plant enzyme (Figure 2 – figure supplement 2), showed a very similar catalytic activity in our hands (Figure 7A, C). It is of note that specific 1-phosphatase activity has been previously reported for the isolated Asp1 phosphatase domain from *Schizosaccharomyces pombe* (Wang et al., 2015). Addition of either magnesium ions or a metal chelator had no detectable effect on VIH2 phosphatase activity during overnight incubation (Figure 7C). We next assayed if the VIH2-PD^RH/AA^ mutations in the phosphatase domain (Figure 7 – figure supplement 7B, C) would abolish the phosphatase activity of the mutant proteins. To our surprise we find VIH2-PD^RH/AA^ to be catalytically active using either 1PP-InsP5 or 5-PP-InsP_7_ as substrate (Figure 7D). We further characterized the kinetic properties of the VIH2-PD^RH/AA^ mutant protein, and found it very similar to wild-type VIH2-PD in time course experiments (Figure 7E, F). Together, our *in vitro* assays of the isolated phosphatase domains suggest that the C-terminus of the bifunctional VIH2 enzyme may be able to hydrolyze different PP-InsP isoforms, but not InsP_6_.

**Figure 7.**
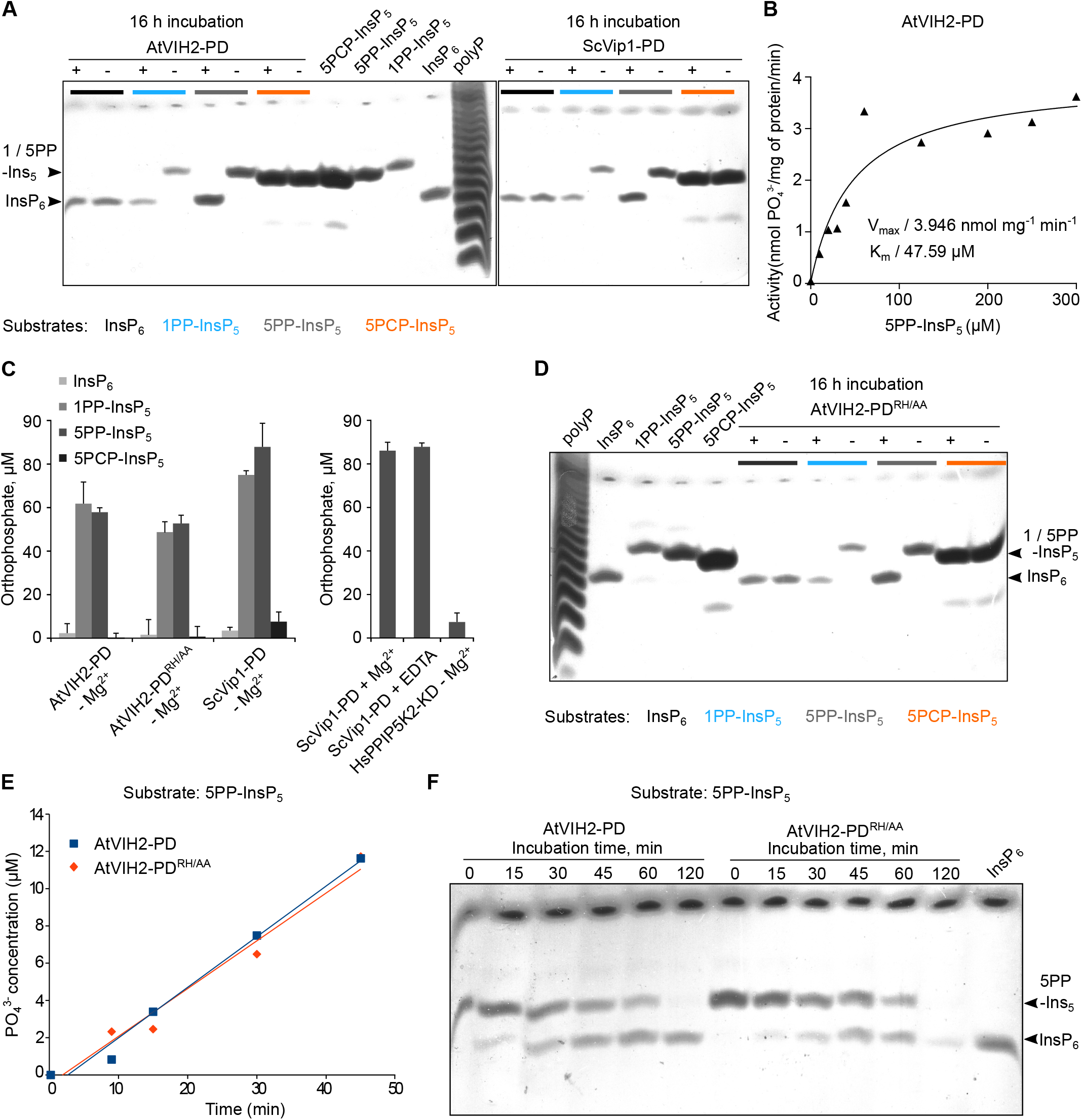
The phosphatase domain of AtVIH2 displays 1 - and 5 - pyrophosphatase activity. **(A)** Reactions containing recombinant phosphatase domain of AtVIH2 (AtVIH2-PD) or ScVip1 (ScVip1-PD) were incubated with 175 μM InsP_6_ (black), 1PP-InsP_5_ (blue), 5PP-InsP_5_ (gray) or a non-hydrolyzable 5PCP-InsP_5_ analog (orange) for 16 h at 37°C. 40 μl of the reaction were separated in a 35% acrylamide gel. The bands corresponding to InsP_6_ and 1 or 1 / 5PP-InsP_5_ are indicated by an arrow head. **(B)** Michaelis-Menten plot of AtVIH2-PD vs. 5PP-InsP_5_. indicate a turnover number of ~0.3/min. **(C)** Phosphatase activity assay. Reactions containing recombinant AtVIH2-KD, AtVIH2-PD^RH/AA^ or ScVip1-PD were incubated with InsP_6_, 1PP-InsP_5_, 5PP-InsP_5_ or 5PCP-InsP_5_ for 16 h at 37°C (left). 1mM Mg^2+^ or 5 mM EDTA were supplemented where indicated (right; 5PP-InsP_5_ only).Recombinant HsVIP2-KD was used as a negative control and tested only with 5PP-InsP_5_ (right). Reactions were performed in quadruplicates and released orthophosphate was quantified using a malachite green assay (Baykov et al., 1988). **(D)** Gel-based PP-InsP phosphatase assay. Reactions were performed with recombinant AtVIH2-PD^RH/AA^ as described in (A), and separated on a 35% acrylamide gel. The bands corresponding to InsP_6_ and 1 or 1 / 5PP-InsP_5_ (stained using toluidine blue) are indicated by arrow heads. **(E)** Time-course experiment comparing the enzymatic activities of recombinant AtVIH2-PD or AtVIH2-PD^RH/AA^. 2 μM enzyme and 40 μM 5PP-InsP_5_ were incubated at 37°C and released orthophosphate was quantified by a malachite green assay. **(F)** Gel-based PP-InsP phosphatase assay. 40 μl of reactions from (E) were separated on a 35% acrylamide gel. The bands corresponding to InsP_6_ and 5PP-InsP_5_ (stained using toluidine blue) are indicated by arrow head.

### PPIP5Ks may respond to changes in cellular ATP levels

Finally, we wanted to investigate whether the PPIP5K kinase and phosphatase domains cooperate in the context of the full-length enzyme. We could express and purify full-length Arabidopsis VIH1 and 2 but found the purified protein to be aggregated in size-exclusion chromatography experiments (Figure 8A). We thus produced the related full-length Vip1 from *S. cerevisiae*, which we could purify to homogeneity and which behaves as a monomer in size-exclusion chromatography (Figure 8B). We next incubated ScVip1 with 5PP-InsP_5_ while varying ATP substrate concentrations. We find that at low ATP concentrations the ScVip1 phosphatase activity predominates, releasing InsP_6_ (Figure 8C). Increasing ATP levels results in stimulation of the PP-InsP kinase activity, producing increasing amounts of InsP_8_. Interestingly, at intermediate ATP concentrations, the kinase and phosphatase activity may be balanced, as we do not observe net production of either InsP6 or InsP_8_ (Figure 8C). Thus, changes in cellular ATP levels may affect PPIP5K kinase and phosphate activities to regulate cellular PP-InsP levels.

**Figure 8.**
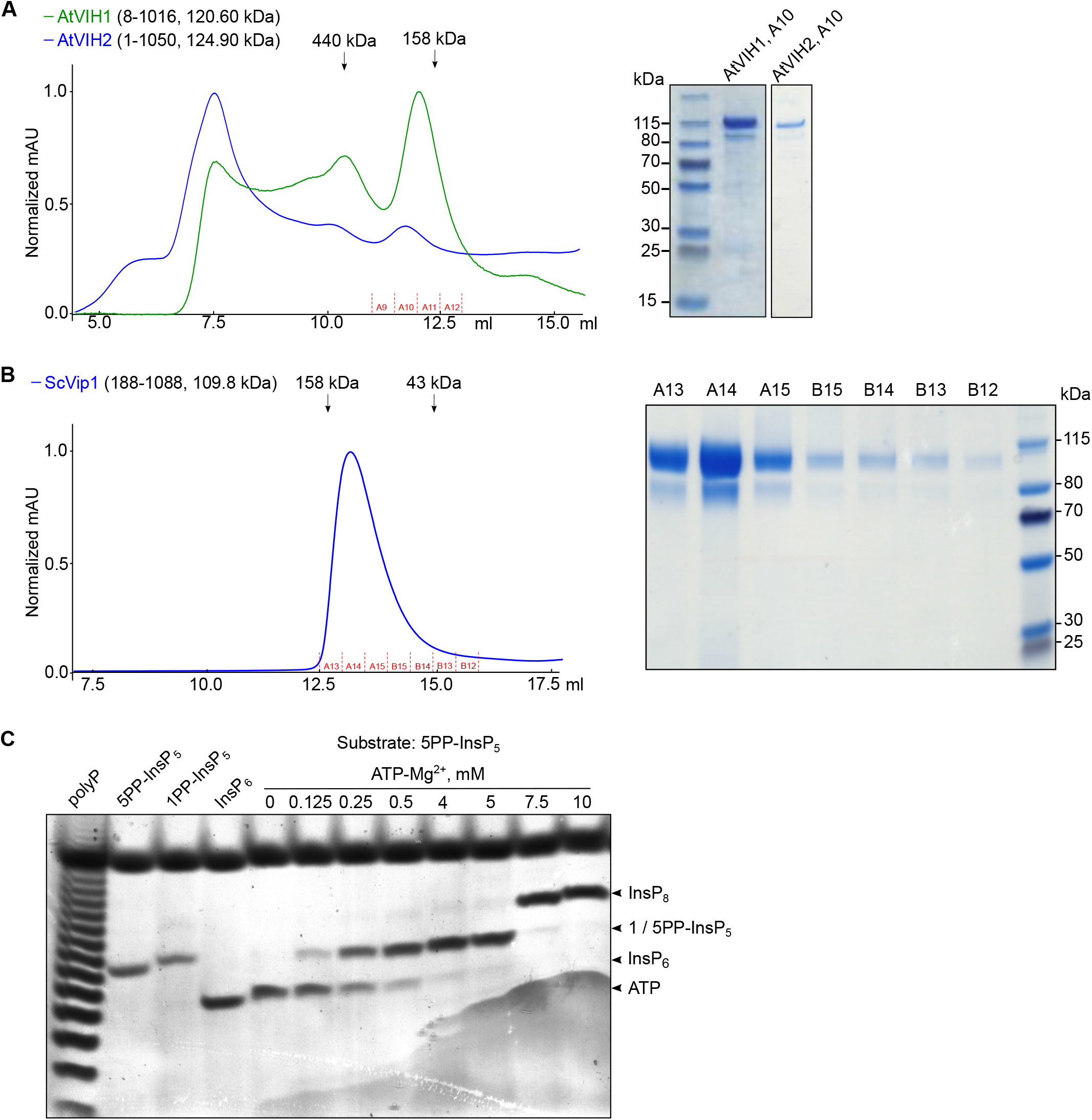
The relative ScVip1 PP-InsP kinase and phosphatase activities change in response to changes in ATP-Mg^2+^ concentrations. **(A)** Size-exclusion chromatography traces of purified full-length AtVIH1 (Gly8 - Thr1016) and AtVIH2 (Met1 – Ser1050). A coomassie-stained SDS-PAGE gel showing the content of the peak fraction (A10) is shown alongside. **(B)** Size-exclusion chromatography trace of purified recombinant ScVip1 (188G-1088R) and fractions harvested for this enzymatic assay shown in C (fractions A13-B12). A coomassie-stained SDA-PAGE gel showing the respective fractions of ScVip1 is shown alongside. **(C)** Bi-functional PP-InsP activity assay of ScVip1. Reactions containing 2 μM protein, 40 μM 5PPInsP_5_ and various various ATP-Mg^2+^ concentrations were incubated at 37°C for 45 min. Product PPInsPs were separated on a native PAGE gel and stained with toluidine bluw. The bands corresponding to InsP_6_, 1 / 5PP-InsP_5_, InsP_8_ and ATP are indicated by arrow heads.

## Discussion

Plants have evolved multiple strategies to maintain cellular Pi homeostasis in Pi deficient conditions. Components involved in this process include PHR family transcription factors, their SPX domain interaction partners, phosphate transporters, ubiquitin ligases controlling phosphate transporter protein levels, microRNAs and ferroxidase enzymes (Puga et al., 2017). The recent finding that eukaryotic SPX domains are cellular receptors for PP-InsPs (Wild et al., 2016) and that PP-InsP binding can regulate the activity of PHR transcription factors (Puga et al., 2014; Wang et al., 2014; Wild et al., 2016; Qi et al., 2017) and phosphate transporters (Wild et al., 2016; Potapenko et al., 2018) prompted us to investigate if the Arabidopsis diphosphoinositol pentakisphosphate kinases VIH1 and 2 are components of the plant phosphate starvation response pathway. Our experiments demonstrate that VIH1 and VIH2 redundantly regulate Pi homeostasis, growth and development. Deletion of the phosphate starvation response transcription factors PHR1 and PHL1 (Rubio et al., 2001; Bustos et al., 2010) can partially rescue the *vih1-2 vih2-4* mutant phenotype. This confirms that VIHs, their PP-InsP reaction products and PHR1/PHL1 are part of a common signaling cascade. Our work is in good agreement with the recent observation that mutations in the inositol polyphosphate biosynthesis pathway impact Pi signaling (Kuo et al., 2018). The fact that our quadruple mutant does not resemble wild-type plants may indicate that PP-InsPs regulate the activity of other components of the phosphate starvation response, such as additional transcription factors (Sun et al., 2015; Wang et al., 2018), phosphate transporters (Liu et al., 2015, 2016; Wild et al., 2016), or other SPX-domain containing proteins (Park et al., 2014). The strong growth and developmental phenotypes of the *vih1-2 vih2-4* double mutant may also hint at additional signaling functions for PP-InsPs unrelated to Pi homeostasis (Tan et al., 2007; Sheard et al., 2010; Mosblech et al., 2011; Laha et al., 2015, 2016; Wild et al., 2016).

It is known that several PP-InsP isoforms are present in eukaryotic cells. Our biochemical analysis of the isolated VIH2 kinase domain reveals that InsP_8_ is the likely reaction product of the VIHs in Arabdiopsis, and thus the biochemical activities of plant, yeast and human PPIP5Ks may be very similar (Mulugu et al., 2007; Pascual-Ortiz et al., 2018; Lin et al., 2009). This hypothesis is supported by our observation that the *vih1-2 vih2-4* mutant can be complemented with the human enzyme carrying mutations in its phosphatase domain (Figure 5I). In line with our biochemical experiments, it has been previously shown that *in vivo* InsP_8_ levels decrease and that InsP_7_ accumulates in *vih2-4* mutant plants (Laha et al., 2015) and that InsP_8_ levels are reduced in PPIP5K1 and 2 double mutants (Gu et al., 2017b). Importantly, the 1PP-InsP5 kinase activity of VIHs is essential for plant growth and development, as a mutations targeting the kinase active site cannot rescue the *vih1-2 vih2-4* mutant phenotypes.

Eukaryotic diphosphoinositol pentakisphosphate kinases are bifunctional enzymes, which in addition to their 1PP-InsP5 kinase domain also harbor a C-terminal phosphatase domain (Shears et al., 2017). For fission yeast Asp1 and human PPIP5K2 it has been previously shown that the phosphatase domain has inositol pyrophosphatase activity, with a preference for C1 pyrophosphorylated substrates (Gu et al., 2017a; Pascual-Ortiz et al., 2018). In our biochemical assays with the Arabidopsis VIH2 and yeast Vip1 phosphatase domains, we observed inositol pyrophosphatase activity against both C1 and C5 pyrophosphorylated substrates (Figure 7A-C).

This suggests that the C-terminal phosphatase domain of diphosphoinositol pentakisphosphate kinases is able to dephosphorylate the reaction product of the kinase domain. Previously, single point mutations targeting the phosphatase domain have been assayed *in vivo* and *in vitro*, with conflicting results (Mulugu et al., 2007; Choi et al., 2007; Wang et al., 2015; Pascual-Ortiz et al., 2018). The phosphatase activity of our VIH2-PD^RH/AA^ mutant protein is virtually indistinguishable from the wild-type enzyme (Figure 7C-F). Given that the catalytic mechanism of histidine acid phosphatase domains in diphosphoinositol pentakisphosphate kinases is currently unknown, we currently cannot generate a phosphatase-dead enzyme for *in planta* analysis. In our VIH2^RH/AA^ complementation lines, we do observe a significant reduction in shoot Pi content, suggesting that the phosphatase domain has a regulatory function in Pi signaling in plants (Figure 3E-H). Our point mutations in the phosphatase domain may either perturb the catalytic activity of the enzyme *in vivo*, or affect the regulatory interplay between the kinase and phosphatase modules. In line with this, our *vih2-6* mutant, which may express a kinase-only VIH2 protein, shows similar phenotypes (Figure 3I-L) (Kuo et al., 2018), and we can rescue our *vih1-2 vih2-4* double mutant only with a phosphatase-mutated version of human PPIP5K2 (Figure 5I-L). Over-expression analysis of kinaseand phosphatase-mutated versions of VIH2 suggest that their enzymatic activities may antagonistically regulate Pi homeostasis (Figure 5A-D). Together, our genetic and biochemical analyses argue for concerted VIH PP-InsP kinase and phosphatase activities in Arabidopsis, which together regulate plant Pi homeostasis, growth and development.

It has been previously proposed that plants may directly sense cellular Pi levels by inorganic phosphate directly binding to SPX sensor domains (Puga et al., 2014; Wang et al., 2014). Our structural and quantitative biochemical characterization of SPX domains as receptors for PP-InsPs together with the genetic evidence presented here now argues for a function of PP-InsPs as central signaling molecules in plant Pi homeostasis and starvation responses. It has been well established in yeast that PP-InsP levels change in response to changes in Pi availability (Lonetti et al., 2011; Wild et al., 2016). How changes in Pi levels *in planta* may relate to accumulation or depletion of PPInsPs is poorly understood (Azevedo and Saiardi, 2017), but our genetic findings now allow us to speculate that the VIH kinase and phosphatase activities may be involved in this process. In this respect it is of note that cellular ATP levels are reduced when extracellular Pi becomes limiting (Choi et al., 2017; Gu et al., 2017a). We thus assayed the kinase and phosphatase activities of full-length Vip1 from yeast, which we can purify to homogeneity from insect cells, in the presence of different ATP concentrations. Indeed, we find that at low ATP-Mg^2+^ levels, Vip1 mainly acts as PPInsP phosphatase, while at higher ATP-Mg^2+^ concentrations significant InsP_8_ synthesis can be observed (Figure 8). We thus speculate that eukaryotic diphosphoinositol pentakisphosphate kinases/phosphatases have evolved to precisely sense cytosolic ATP concentration, or ATP/ADP ratios, relating changes in Pi levels to changes in PP-InsP pools. Our genetic data support a signaling cascade in which specific PP-InsP isoforms, possibly 1,5-(PP)2-InsP_4_, in turn trigger Pi starvation responses or maintain Pi homeostasis by interacting with their SPX receptors being present in many plant phosphate signaling proteins. Future studies will be required to test this hypothesis in biochemical, mechanistic and physiological terms.

## Materials and Methods

### Plant material and growth conditions

Plants were grown at 21°C, 50 % humidity and in a 16 h light : 8 h dark cycle. For root imaging, western blots and RT-PCR, seedlings were grown on vertical plates containing half-strength Murashige and Skoog (^1/2^MS, Duchefa) media containing 1 (w/v) % sucrose, and 0.8 (w/v) % agar (Duchefa) for 6 to 10 d. Seeds were sequentially surface-sterilized using 70 % ethanol, a 5 % hypochlorite solution (Javel, 13 - 14 %), and rinsed 4 times with sterilized water.

The Arabidopsis seeds of T-DNA insertion lines were obtained from the European Arabidopsis Stock Center (http://arabidopsis.info/) except for *phr1 phl1*, *ipk1-1*, which were kindly provided by Dr. Javier Paz-Ares (National Center for Biotechnology, Madrid, Spain) and Dr. Yves Poirier (University of Lausanne, Switzerland). The T-DNA lines used in this study are as follows: *vih1-2* (SALK_094780C), *vih1-4* (WiscDsLox293-296invF12), *vih1-6* (SAIL_175_H09), *vih2-4* (GK_080A07), *vih2-6* (GK_008H11), *spx1-1* (SALK_092030C), *spx2-1* (SALK_080503C), *phl1* (SAIL_731_B09), *ipk1-1* (SALK_065337). *phr1* used in this study is an EMS mutant (Rubio et al., 2001).

### Characterization of T-DNA mutants

Homozygous lines were identified by PCR using T-DNA left and right border primers paired with gene-specific sense and antisense primers (Supplementary Table 2A). mRNA expression levels were quantified using cDNA prepared from RNA extracts of seedlings 9 d after germination (DAG). Primers flanking the T-DNA position of the *vih1-2* allele were used in qRT-PCR quantifications of VIH1 expression for *vih1-2*, *vih1-4* and *vih1-6*. VIH2 expression in *vih2-4* and *vih2-6* plants was quantified using primers flanking the respective T-DNA positions. Primer sequences are listed in Supplementary Table 2D. The crosses between different T-DNA lines and the analysis of their segregation ratios are listed in Supplementary Table 3A, B.

### Generation of transgenic lines

All transgenic Arabidopsis lines are listed in Supplementary Table 1. VIH1 and VIH2 and SPX1 and 3 were amplified from Arabidopsis cDNA and introduced into either pB7m34GW or pH7m34GW binary vectors. Mutations targeting the kinase or phosphatase domains were introduced by site-directed mutagenesis (mutagenesis primers listed in Supplementary Table 2B). VIH1 targeting amiRNAs were designed using WMD3 – Web MicroRNA Designer (http://wmd3.weigelworld.org/). Fragments containing AtVIH1^amiRNA^ were amplified and introduced into the pMDC7 binary vector using the primers listed in Supplementary Table 2B (Curtis and Grossniklaus, 2003). Fragments coding for full-length HsVIP2, its kinase (HsVIP2-KD) or its phosphatase (HsVIP2PD) domain were amplified by PCR (primers listed in Supplementary Table 2B) using a synthetic HsVIP2 gene (codon was optimized for expressing in Arabidopsis, Geneart-ThemoFisher) as template.

Binary vectors were assembled by the multi-site Gateway technology (Thermo Fisher Scientific). All constructs were transformed into *Agrobacterium tumefaciens* strain pGV2260. Plants were transformed using the floral dip method (Clough and Bent, 1998). Transformants were selected in ^1/2^MS medium supplemented with BASTA or Hygromycin down to T3 generation.

### Inorganic Pi concentration determination

To measure inorganic Pi concentration at seedling stage, seedling 7 DAG were transferred from vertical plates containing ^1/2^MS, 1 (w/v) % sucrose, and 0.8 (w/v) % agar to new vertical plates containing Pi deficient ^1/2^MS (Duchefa), 1 (w/v) % sucrose, and 0.8 (w/v) % agarose (A9045, Sigma), and supplemented with varying concentrations of Pi (KH_2_PO_4_/K_2_HPO_4_ mixture, pH 5.7). To measure inorganic Pi concentration at adult stage, seedlings 7 DAG were transplanted from vertical plates to soil. Seedlings 14 or 19 DAG and adult plants 20 DAG were dissected and shoot/root Pi content was measured by the colorimetric molybdate assay (Ames, 1966).

### Plant protein extraction for western blot

Around 250 mg seedlings 10 DAG was harvested and frozen in liquid nitrogen in 2 ml eppendorf tubes with metal beads and ground in a tissue lyzer (MM400, Retsch). Ground plant material was resuspended in 500 μl extraction buffer (150 mM NaCl, 50 mM Tris-HCl pH 7.5, 10 % (v/v) glycerol, 1 % (v/v) Triton X-100, 5 mM DTT) containing a protease inhibitor cocktail (P9599, Sigma, 1 tablet/20 ml), centrifuged for 30 min at 4,000 rpm and at 4°C. Protein concentrations were estimated using a Bradford protein assay. Around 100 μg protein was mixed with 2x SDS loading buffer, boiled at 95°C for 10 min and separated on 8 % SDS-PAGE gels. Anti-GFP antibody coupled with horse radish peroxidase (HRP, Miltenyi Biotec) at 1:2000 dilution was used to detect eGFP/mCit tagged protein constructs.

### RNA analyses

Around 200 mg seedlings 10 DAG was harvested in 2 ml eppendorf tubes and frozen in liquid nitrogen. 1-2 μg total RNA extracted using a RNeasy plant mini kit (Qiagen, Valencia, CA) was reverse-transcribed using the SuperScript VILO cDNA synthesis kit (Invitrogen, Grand Island, NY). qRT-PCR amplifications were monitored using SYBR-Green fluorescent stain (Applied Biosystems), and measured using a 7900HT Fast Real Time PCR-System from Applied Biosystems (Carlsbad, CA). For the normalization of gene transcripts, *ACTIN2* or *ACTIN8* was used as a internal control (primers listed in Supplementary Table 2D). Averages of triplicate or quadruplicate reactions ± SE are shown. All of the quantifications were repeated at least 3 times, with similar results.

### Confocal laser scanning

Zeiss LSM780 equipment was used for confocal laser scanning. Confocal settings were set to record the emission of eGFP (excitation 488 nm, emission 500–550 nm), and mCit (excitation 514 nm, emission 524–580 nm). For the imaging of Arabidopsis root tips and pollen, a 40 x / 1.3 oil objective and 20 x / 0.50 objective were used, respectively. Images were acquired with Zen 2012 SP2 using identical settings for all samples. All image analyses were performed in the FIJI distribution of ImageJ 1.51w (http://imagej.nih.gov/ij/).

### GUS staining

Fragments encompassing 1.85 kb and 1.80 kb of the promoter regions of VIH1 and VIH2, respectively, were amplified from Arabidopsis genomic DNA by using primers listed in Supplementary Table 2B, and cloned using the Gateway LR reaction to generate pB7m34GW binary vectors expressing pVIH1::GUS and pVIH2::GUS. Plant tissues from the transgenic lines were fixed in sodium phosphate buffer (50 mM sodium phosphate, pH 7) containing 2 (v/v) % formaldehyde for 30 min at room temperature and subsequently, washed 2 times with the sodium phosphate buffer. The fixed plant tissues were suspended in staining buffer (50 mM sodium phosphate, 0.5 mM K - ferrocyanide, 0.5 mM K - ferrocyanide, 1 mM X-GlcA) vacuumed infiltrated for 15 min, and stained for 4-6 h at 37°C. The stained plant tissues were then imaged directly or stored in 20 % EtOH after two washes with 96 and 60 (v/v) % ethanol, respectively, for 1 h. Images were acquired with a Zeiss StiREO Discovery.V8 microscope.

### Protein expression and purification

Synthetic genes coding for AtVIH1 (residues 1 - 1049), AtVIH2 (residues 1 - 1050), HsVIP2 (residues 1 - 1124) and ScVip1 (residues 1 - 1146), codon optimized for expression in *Spodoptera frugiperda* Sf9 cells were ordered from Geneart (ThemoFisher). The fragments coding for the kinase domain of AtVIH2 (residues 11 - 338), HsVIP2 (residues 41 - 366), the fragments coding phosphatase domain of AtVIH2 (residues 359 - 1002), ScVIP2 (residues 515 - 1088), and the fragments coding both the kinase domain and the phosphatase domain of AtVIH1 (residues 8 - 1016), AtVIH2 (residues 1 - 1050) and ScVip1 (residues 188 - 1088) were amplified from the synthetic genes. Point mutations were introduced by site-directed mutagenesis. All DNA fragments were cloned into a modified pFastBac1 vector (Geneva Biotech), providing a N-terminal TEV (tobacco etch virus protease) cleavable Strep-9xHis tandem affinity tag (primers are listed in Supplementary Table 2C), and all constructs were confirmed by DNA sequencing.

Bacmids were generated by transforming the plasmids into *Escherichia coli* DH10MultiBac (Geneva Biotech), isolated, and transfected into *Spodoptera frugiperda* Sf9 cells with profectin transfection reagent (AB Vector) followed by viral amplification. All the proteins were expressed in *Spodoptera frugiperda* Sf9 cells and harvested from the medium 3 days post infection, and purified separately by sequential Ni^2+^ (HisTrap excel HP, GE Healthcare) and Strep (Strep-Tactin XT Superflow, IBA, Germany) affinity chromatography. Next, the proteins were further purified by size-exclusion chromatography on a HiLoad 26/600 Superdex 200pg column (GE Healthcare) or on a Superdex 200 increase 10/300 GL column (GE Healthcare), equilibrated in 150 mM NaCl, 20 mM HEPES pH 7.0. Protein purity and molecular weight were analyzed by SDS - PAGE and MALDITOF mass spectrometry.

### NMR kinase assay

Proteins encoding full-length proteins of AtVIH1 and ScVip1, and kinase-domains of AtVIH2 and HsVIP2 were used. 1 – 10 μM of proteins were incubated in a buffer containing (final concentrations) 20 mM HEPES pH 7, 50 mM NaCl, 1mM DTT, 2.5 mM ATP, 5 mM creatine phosphate, 1 U creatine kinase, 7.5 mM MgCl_2_ and 175 uM of [^13^C_6_]InsP_6_ or [^13^C_6_]5PP-InsP_5_, at a final volume of 550 μl. The reaction was incubated at 37°C overnight, quenched with 50 μl 0.7 M EDTA, lyophilized and resuspended in 600 uL of 100% D_2_O. The samples were measured on an AV 600 from Bruker using standard pulse programs (Harmel *et al*., *in prep*). Kinetic assays were performed similarly with the exception that the labeled InsPs were added just before NMR measurements and were not quenched by EDTA.

### Phosphatase specificity and kinetic assays

Substrates: InsP_6_ was purchased from SiChem, Germany. 1PP-InsP_5_ (Capolicchio et al., 2013); 5PPInsP_5_ (Harmel *et al*., *in prep*), and the non-hydrolyzable 5-methylene-bisphosphonate inositol pentakisphosphate (5PCP-InsP_5_) (Wu et al., 2013) were synthesized.

Purified recombinant AtVIH2-PD (8.3 µg), AtVIH2-PD^RH/AA^ (5µg), ScVip1-PD (27 µg) and HsVIP2-KD (17.3 µg) were incubated with 175 µM of each inositol polyphosphate substrate for 16 h at 37°C in 150 mM NaCl, 20 mM HEPES pH7, 1 mg/ml BSA. The reactions were stopped by freezing in liquid nitrogen. Reaction products were separated by electrophoresis in 35 % acrylamide gels in TBE buffer (89 mM Tris, pH8, 89 mM boric acid, 2 mM EDTA) and stained with toluidine blue (20% methanol, 0.1% toluidine blue, and 3% glycerol) as described (Losito et al., 2009) Phosphatase activity were measured using the malachite green phosphate assay (Baykov et al., 1988) kit (Sigma), following the manufacturer’s instructions.

### Bifunctional kinase and phosphatase activity assays

2 µM recombinant ScVip1(188 - 1088) was incubated with 80 µM 5PP-InsP_5_ for 16 h at 37°C in a buffer containing 150 mM NaCl, 20 mM HEPEs, pH 7.0, and 1 mg / ml BSA. Variable concentrations of ATP-Mg^2+^ was supplemented into the buffer. The reactions were stopped by freezing in liquid nitrogen, then separated in 35 % acrylamide gels in TBE buffer and stained with toluidine blue.

### Statistics

Simultaneous inference was used throughout to limit the false positive decision rate in these randomized one- or two-way layouts. Designs in which single plants represent the experimental unit were analyzed using a linear model, whereas for designs with technical replicates, a mixed effect model was used accordingly. Normal distributed variance homogeneous errors were assumed when appropriate, otherwise a modified variance estimator allowing group-specific variances was used (Herberich et al., 2010). Multiple comparisons of several genotypes vs. wild-type (Col-0) were performed according to Dunnett (Dunnett, 1955) (Figure 2D; Figure 3C, G, Figure 5K). A Tukey-type all-pairs comparison between the genotypes (either unpooled or pooled, indicated in the respective figure panel by a horizontal line) was used in Figures 2 – figure supplement 5B, Figure 4E, Figure 5C). Trend analyses were performed for qualitative levels by a multiple contrasts test (Williams, 1971) in Figures 2E, F, 4B, and for quantitative levels a regression-type test was empoyed (Tukey et al., 1985) (Figure 4C). Genotype-by-concentration interaction analyses were performed by interaction contrasts (Kitsche and Hothorn, 2014) (Figure 2F). All p-values are for 2-sided tests (*** p < 0.0001; ** p < 0.001; * p < 0.01 for mutant vs. wild-type; ### p < 0.0001; ## p < 0.001; # p < 0.01 for mutant vs. mutant; &&& p < 0.0001; && p < 0.001; & p < 0.01 for mutant vs. alternative mutant). All computations were performed in R version 3.3.2., using the packages multcomp (Hothorn et al., 2008), nlme (Debroy, 2006), tukeytrend (Tukey et al., 1985).

## Acknowledgments

This work was supported by European Research Council under the European Union’s Seventh Framework Programme (FP/2007-2013)/ERC Grant Agreement 310856 (to MH), by Swiss National Foundation Sinergia Grant CRSII5_170925 (to DF and MH) and by an HHMI International Research Scholar Award (to MH). KL was supported by an EMBO long-term fellowship (ALTF-493-2015). RKH and RP were supported by the Leibniz-Gemeinschaft (SAW-2017-FMP-1). We thank D. Couto, L. Lorenzo-Orts, M. Ried and Y. Poirier for critically reading the manuscript.

---

**Figure 2 – figure supplement 1:**
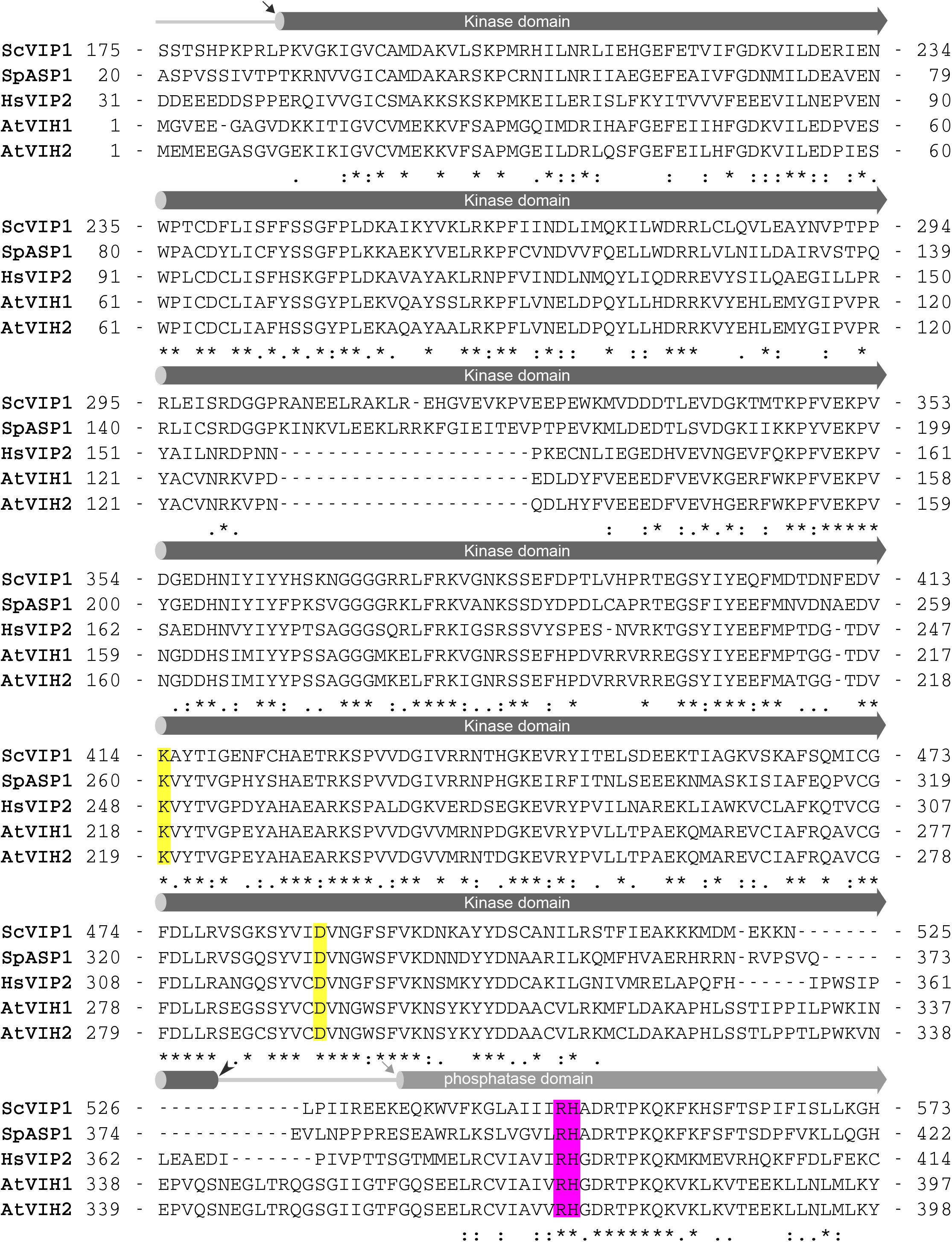
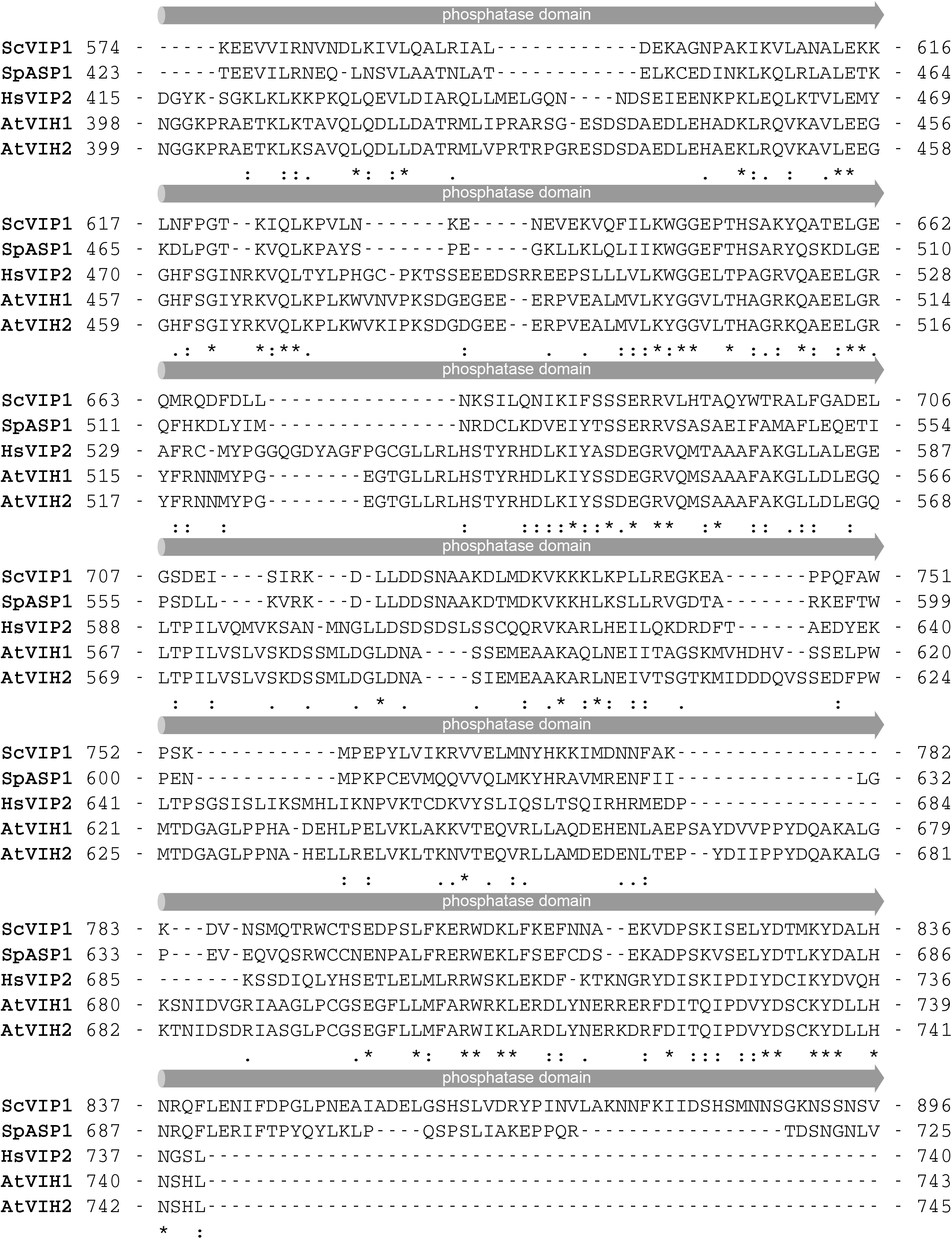
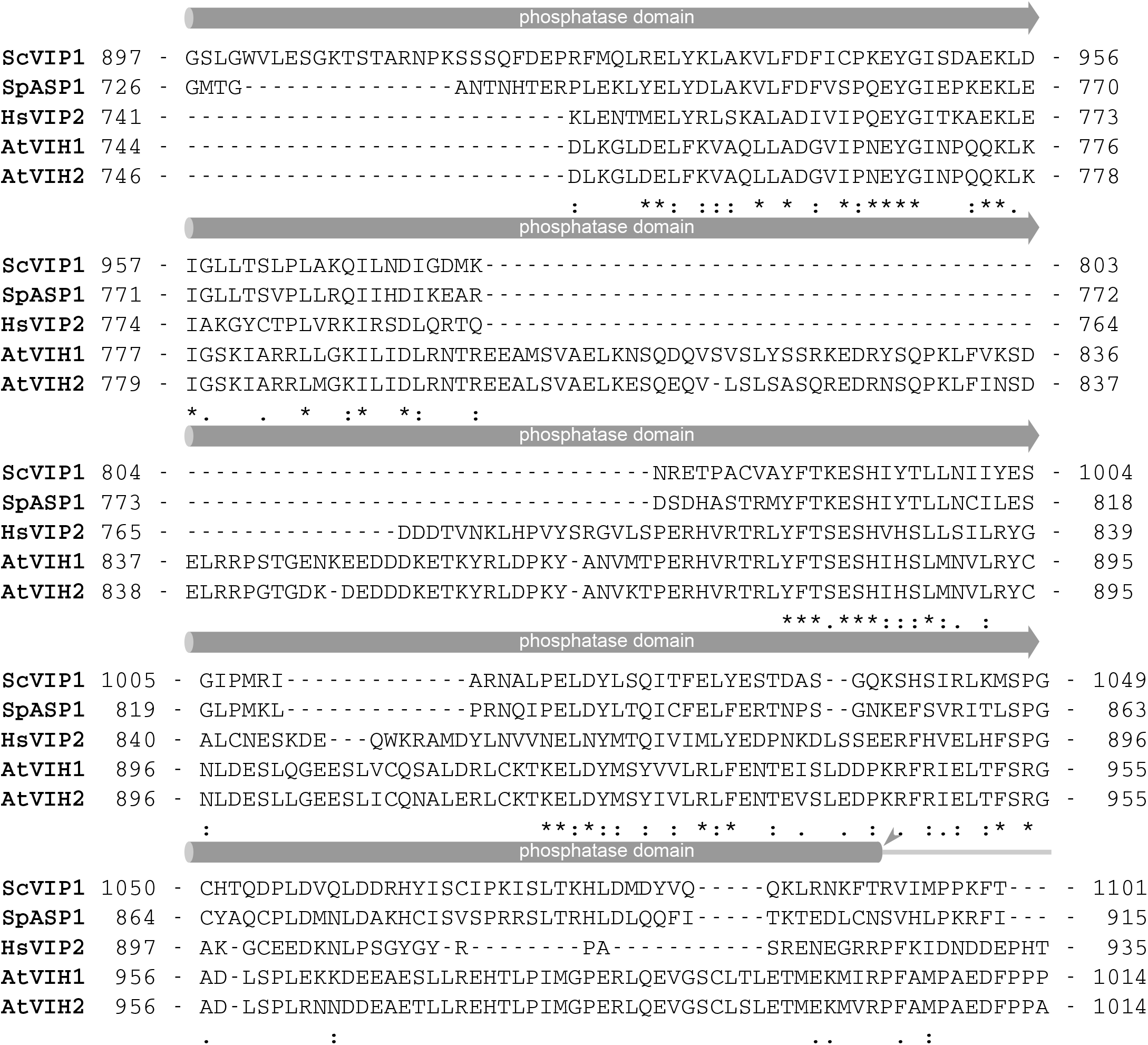
The kinase domain and phosphatase domain of VIHs/PPIP5Ks are conserved among different species. Structure-based sequence alignment of the kinase - phosphatase cores of *Saccharomyces cerevisiae* Vip1 (Uniprot (ttp://www.uniprot.org) identifier: Q06685), *Schizosaccharomyces pombe* ASP1 (Uniprot identifier: O74429), *Homo sapiens* PPIP5K22 (Uniprot identifier: O43314), *Arabidopsis thaliana* VIH1 (Uniprot identifier: Q84WW3), and VIH2 (Uniprot identifier: F4J8C6). Predicted kinase domain and phosphatase domains in VIHs/PPIP5Ks are shown as dark gray and gray cylinders, respectively. Linker regions are indicated by gray lines. Conserved residues that are mutated in this study are highlighted in yellow for the kinase domain and in magenta for the phosphatase domain.

**Figure 2 – figure supplement 2:**
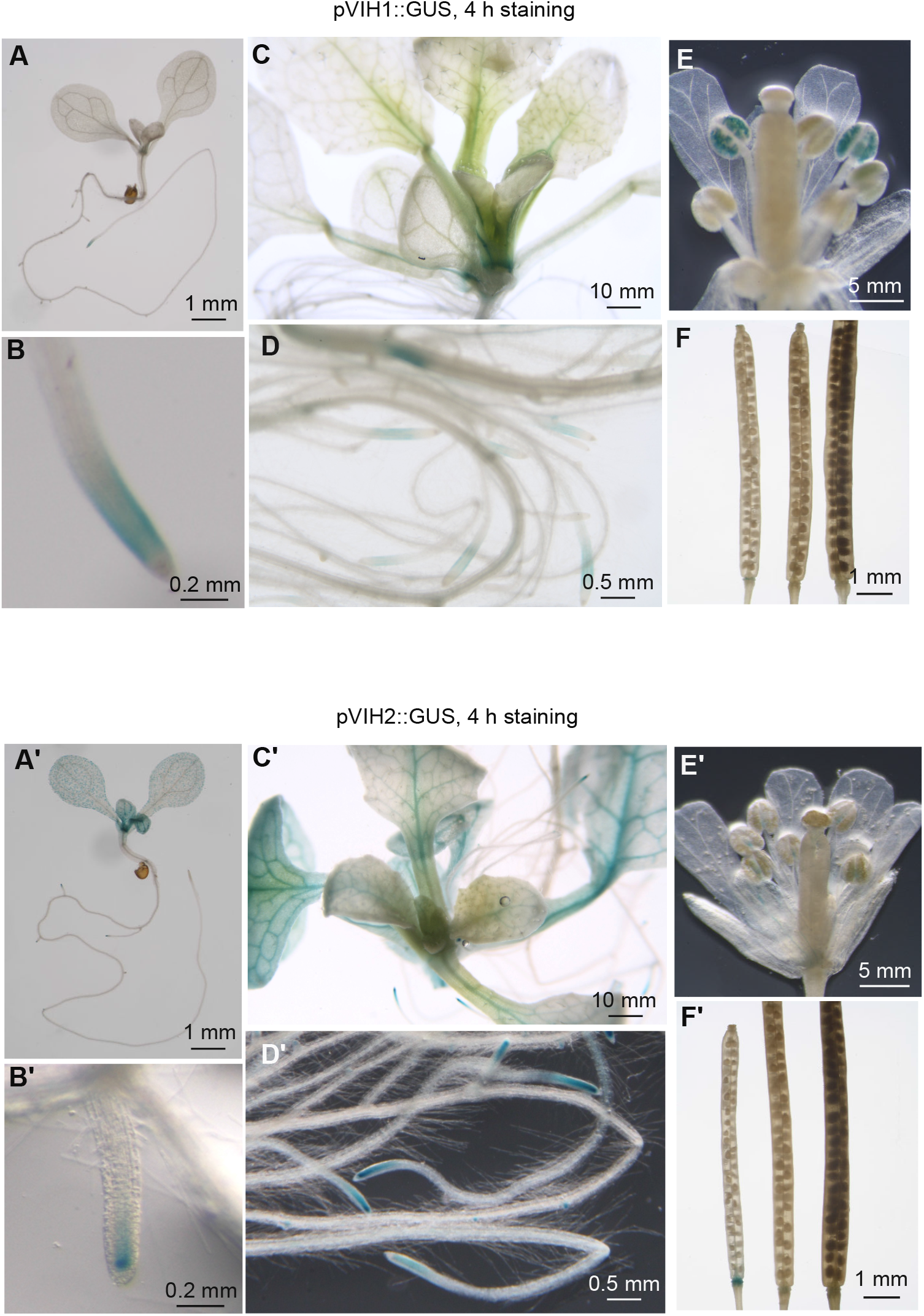
VIH1 and VIH2 show partly unique and partially overlapping expression patterns. Transgenic Arabidopsis lines expressing pVIH1::GUS and pVIH1::GUS in the Col-0 wild-type background were stained for 4 h for β-glucuronidase activity. Images were acquired from seedlings 7 DAG (A,B and A’,B’) and 14 DAG (C,D and C’,D’), flowers (E and E’) and siliques (F and F’).

**Figure 2 – figure supplement 3:**
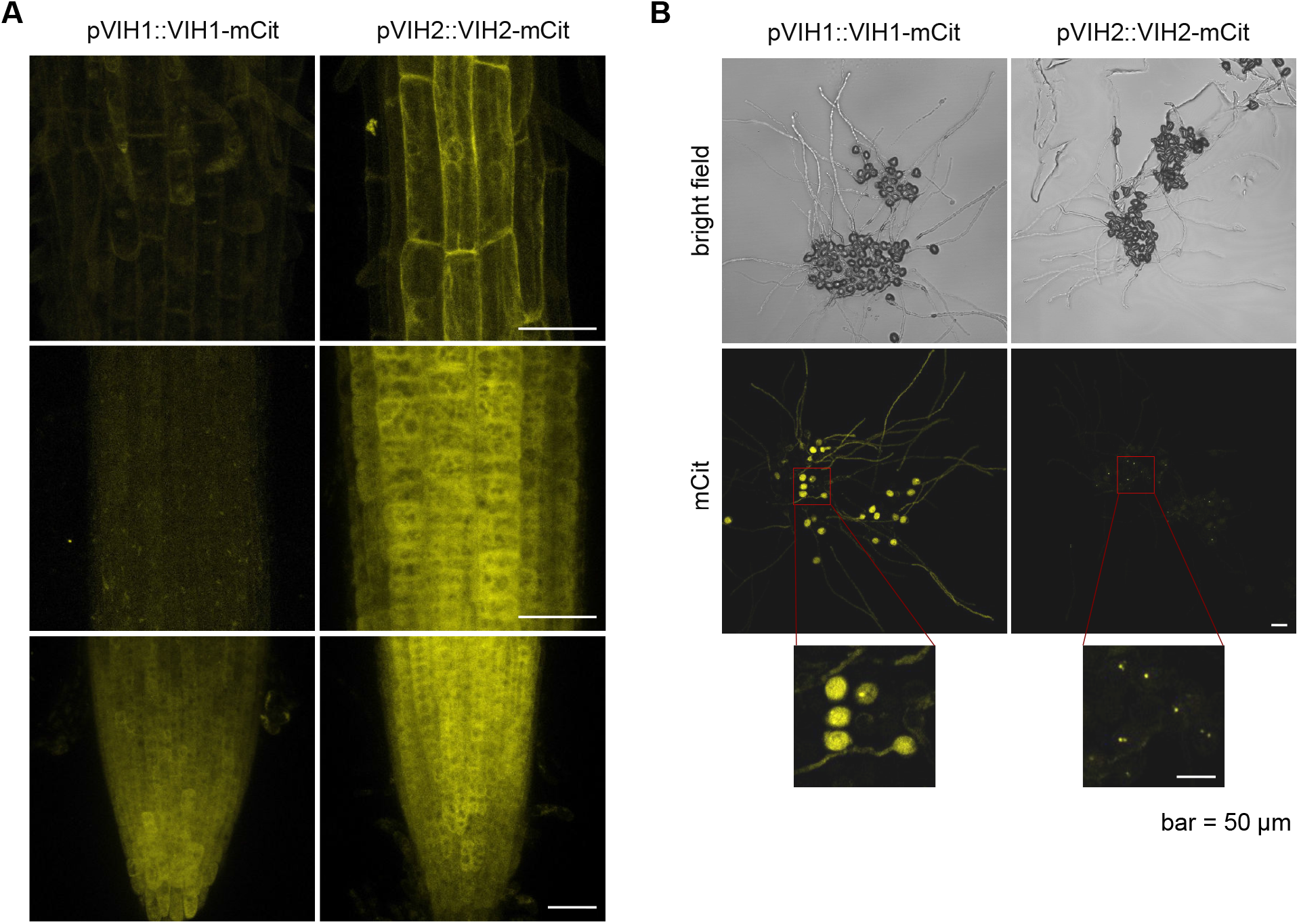
VIH1 and VIH2 are cytoplasmic proteins with partially overlapping expression domains. Stacked slices of (**A**) confocal scanning microscopy images showing the root tip of transgenic Arabidopsis expressing pVIH1::VIH1-mCit (left) and pVIH2::VIH2-mCit (right) in the *vih1-2* and *vih2-4* background, respectively. (**B**) Confocal scanning microscopy images of pollen and pollen tubes from the plants shown in (A), incubated for 4 h on agarose plates containing 5 mM CaCl2, 5 mM KCl, 1 mM MgSO_4_, 0.01 % (w/v) H_3_BO_4_ and 10 % (w/v) sucrose, and 1.5 % (w/v) agarose pH 7.5. Similar localization and expression patterns were observed in at least 3 independent transgenic lines. Scale bars = 50 μm.

**Figure 2 – figure supplement 4:**
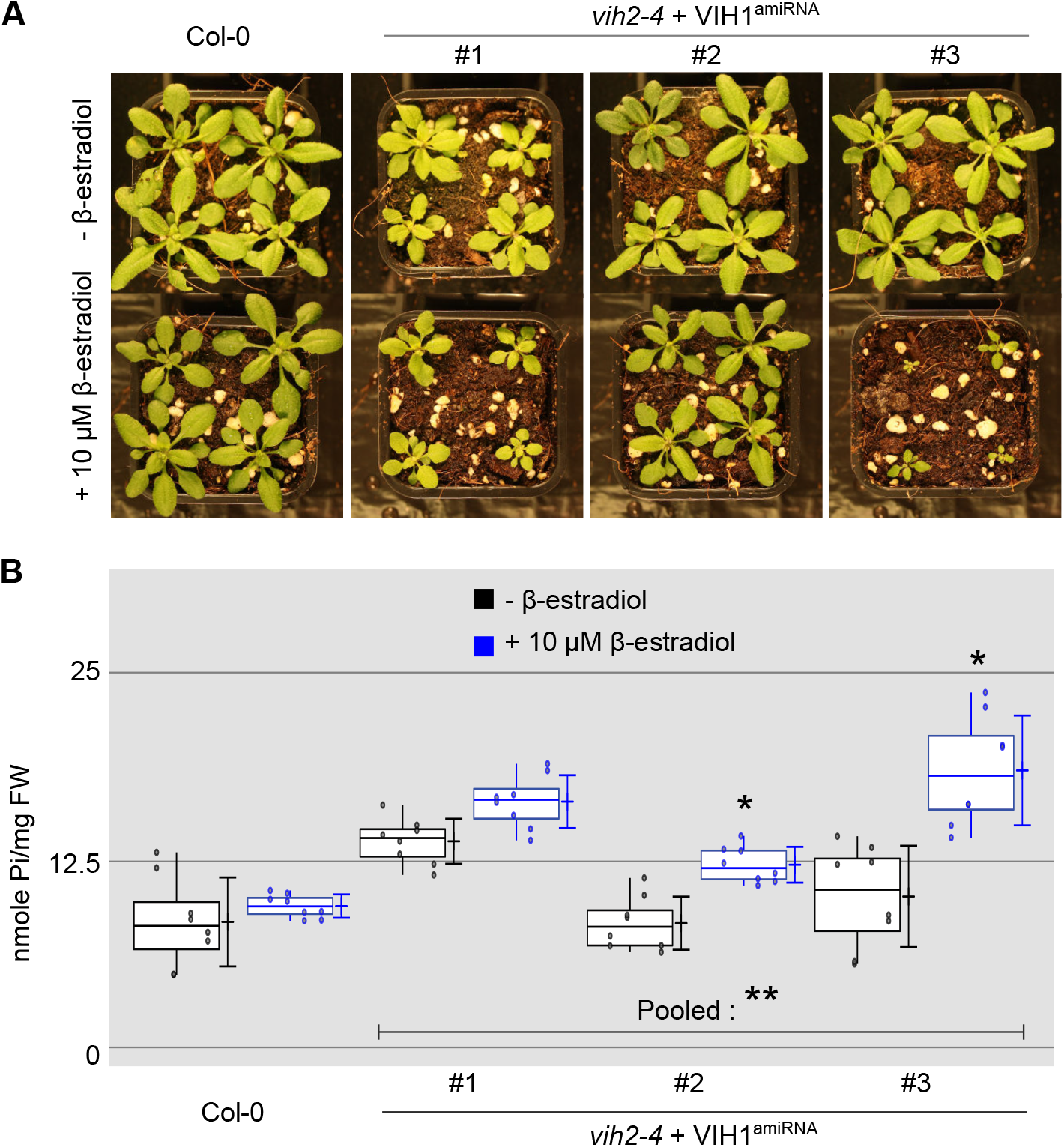
Inducible knock-down of VIH1 in the *vih2-4* background impairs plant growth and leads to shoot Pi accumulation. **(A)** Growth phenotypes of Col-0 wild-type and inducible mutant lines. Seedlings 7 DAG were transferred to soil and sprayed daily with tap water containing either 10 μM β-estradiol diluted from a 100 mM stock solution (100 % [v/v] ethanol), or the corresponding concentration of ethanol only. **(B)** Shoot Pi content of plants 20 DAG treated or not with 10 μM β-estradiol as described in (A). For each boxed position, 4 independent plants were measured using 2 technical replicates.

**Figure 3 – figure supplement 1:**
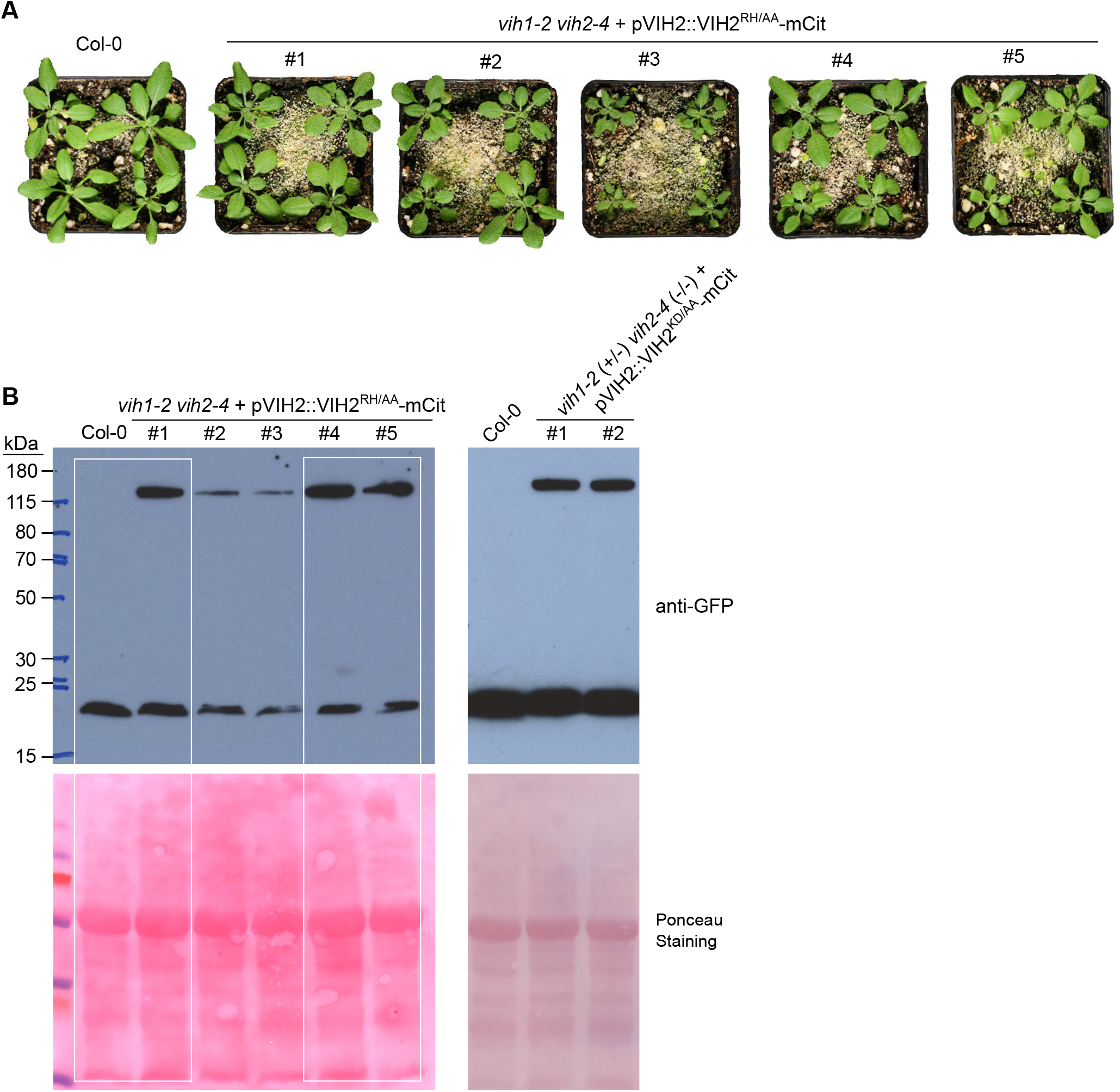
Phenotypes of *vih1-2 vih2-4* double mutant lines complemented with pVIH2::VIH2^RH/AA^-mCit or pVIH2::VIH2^KD/AA^-mCit. **(A)** Shown are growth phenotypes of five independent *vih1-2 vih2-4* lines, complemented with pVIH2::VIH2^RH/AA^-mCit. All plants were transferred to soil 7 DAG and grown for 21 d, Col-0 wild-type plants are shown alongside. **(B)** Left panel shows full western blots of the VIH2^RH/AA^-mCit complementation lines shown in Figure 3E (white boxes); right panel shows western blots of VIH2^KD/AA^-mCit complementation lines which are heterozygous for *VIH1*

**Figure 3 – figure supplement 2:**
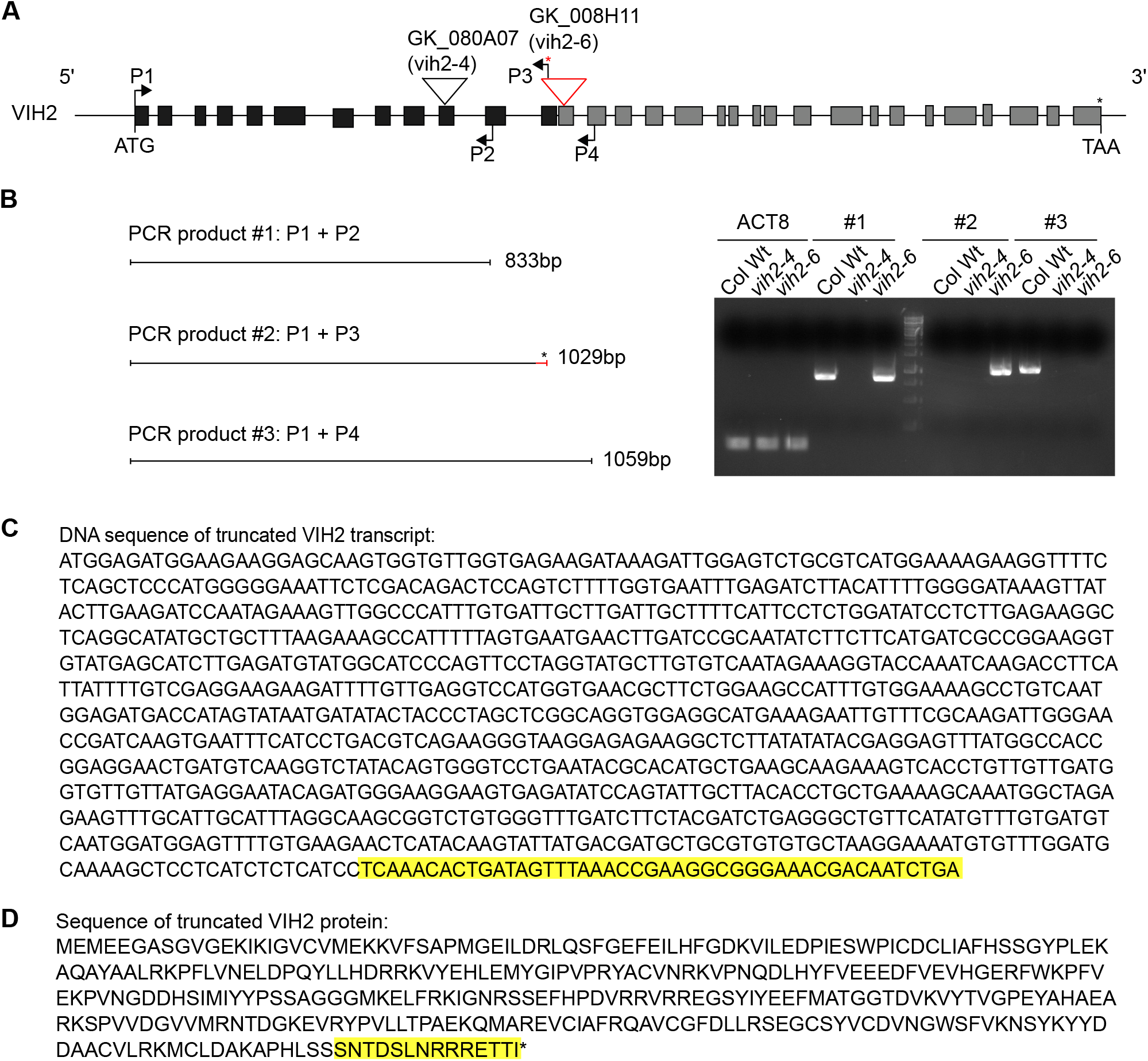
A truncated transcript in the *vih2-6* mutant encodes a truncated VIH2 protein harboring the kinase domain only. **(A)** Insertion positions of T-DNA in *vih2-4* and *vih2-6* alleles and transcript detection by RT-PCR analysis at the At3g01310 locus (*VIH2*). A red asterisk indicates the putative stop codon residing in the T-DNA sequence in the *vih2-6* mutant, which is included in the reverse primer P3 (vih2-*6*_TDNA_R). The primer set P1 (VIH2cDNA_F) + P4 (VIH2-c-R) encompasses the transcript region from the ATG start codon to sequences adjacent to the T-DNA insertion position in *vih2-6*. **(B)** RT - PCR products #1, 2 and 3 are amplified by primer set P1 + P2 (VIH2-b-R), P1 + P3 and P1 + P4 using cDNA reverse-transcribed from 10 DAG Col-0 wild-type, *vih2-4* and *vih2-6* mutant seedlings. Transcripts of ACTIN8 (ACT8) were used as a cDNA input control. **(C)** DNA sequence of the truncated VIH2 transcript in *vih2-6*. The highlighted (yellow) sequence is from the left border of the inserted T-DNA. **(D)** Protein sequence translated from the truncated VIH2 transcript in *vih2-6*. The additional residues (highlighted in yellow) originate from the left border of the T-DNA insertion.

**Figure 5 – figure supplement 1:**
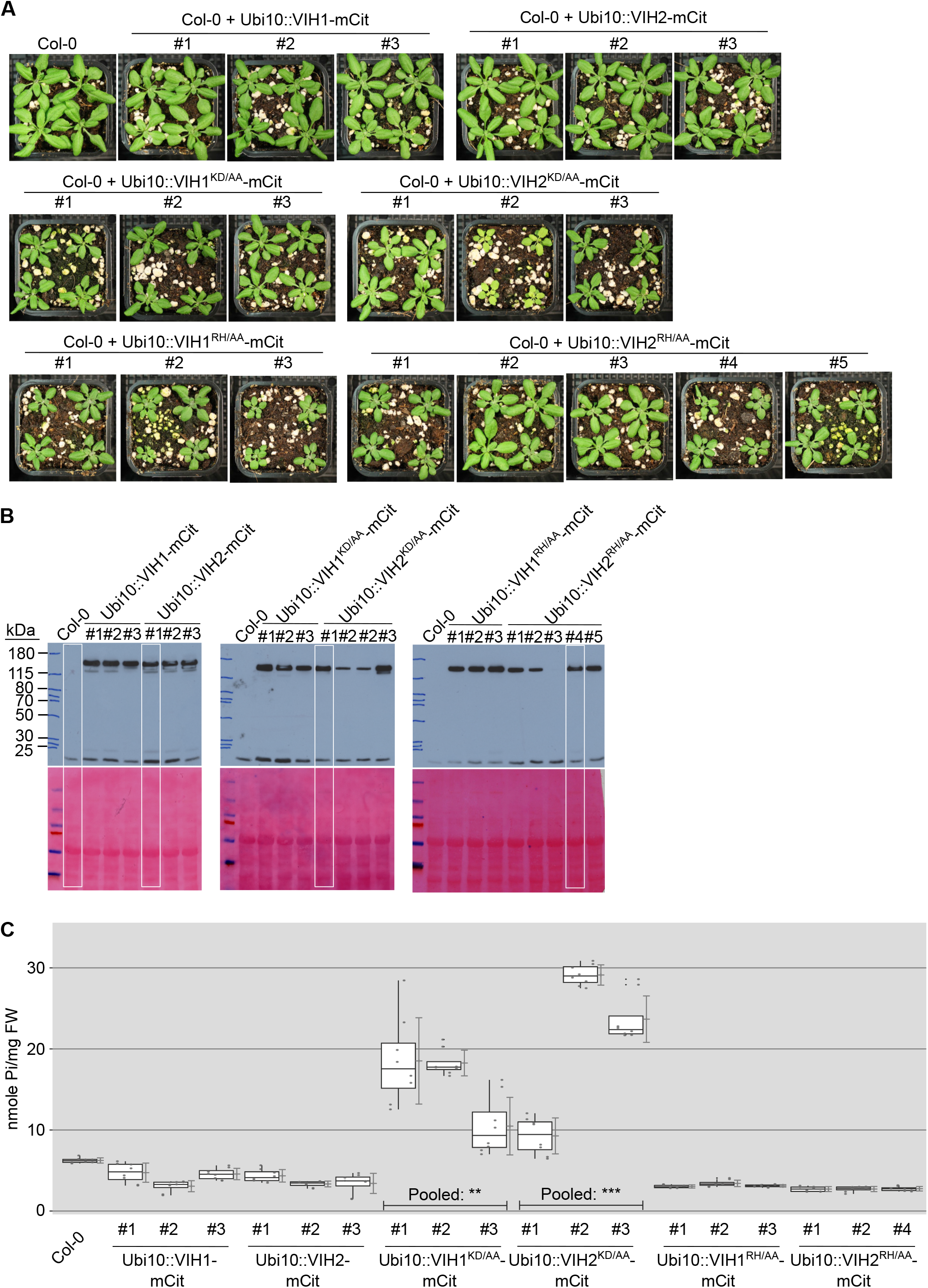
Over-expression of mCit-tagged VIH1 VIH2, and the corresponding kinase and phosphatase domain mutants. **(A)** Growth phenotype of plants expressing Ubi10::VIH1-mCit, Ubi10::VIH2-mCit, Ubi10::VIH1^KD/AA^-mCit, Ubi10::VIH2^KD/AA^-mCit, Ubi10::VIH1^RH/AA^-mCit and Ubi10::VIH2^RH/AA-^mCit. Seedings 7 DAG were transferred to soil and grown for 21 d. **(B)** Expression of VIH1-mCit, VIH2-mCit, VIH1^KD/AA^-mCit, VIH2^KD/AA^-mCit, VIH1^RH/AA^-mCit and VIH2^RH/AA^-mCit proteins as detected by an anti-GFP antibody. White box shows the selected lines for Fig. 5B. **(C)** Shoot Pi contents of plants 20 DAG from selected lines as described in (A). 4 independent plants were measured with 2 technical replicates each.

**Figure 5 – figure supplement 2:**
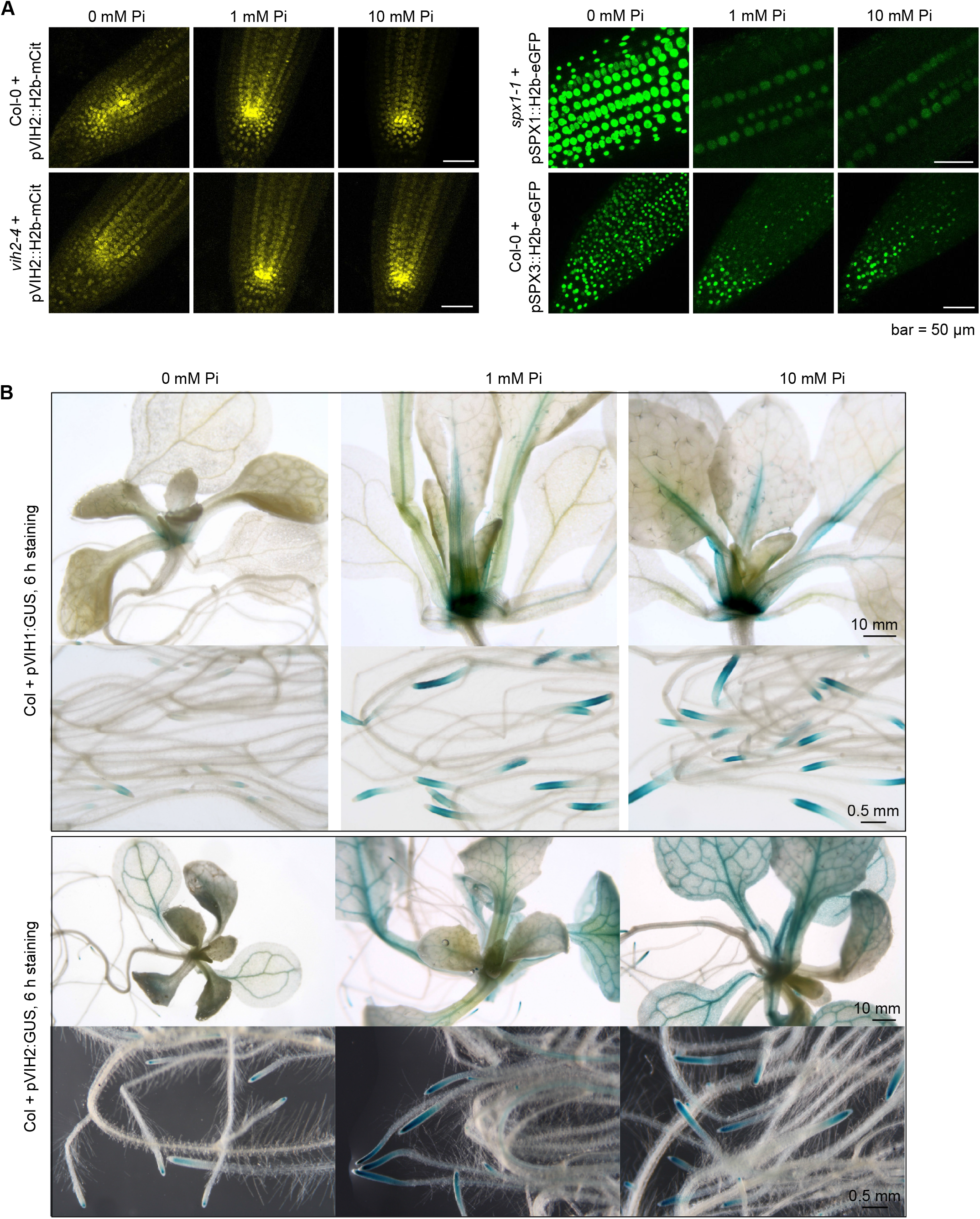
The VIH1 and VIH2 promoters are not induced by low or high Pi treatment. **(A)** Yellow colored representative confocal images showing the expression of mCit-tagged histone H2b under the control of VIH2 promoter in root tip of Col-0 wild-type and *vih2-4* seedlings. Green colored images showing the expression of eGFP-tagged H2b under the control of SPX1 promoter and SPX3 promoter in root tip of *spx1-1* and Col-0 wild-type seedlings, respectively. Seedlings 7 DAG were transferred from ^1/2^MS plates to Pi-deficient ^1/2^MS liquid medium supplemented with 0 mM, 1 mM or 10 mM Pi, and grown for 3 d. Scale bar = 50 μm. **(B)** The staining (6 h) of GUS reporter lines expressing pVIH1::GUS and pVIH2::GUS in a Col-0 wild-type background. Seedlings 7 DAG were transferred from ^1/2^MS plates to Pi-missing ^1/2^MS liquid medium supplemented with 0 mM, 1 mM or 10 mM Pi, and grown for 6 d.

**Figure 6 – figure supplement 1:**
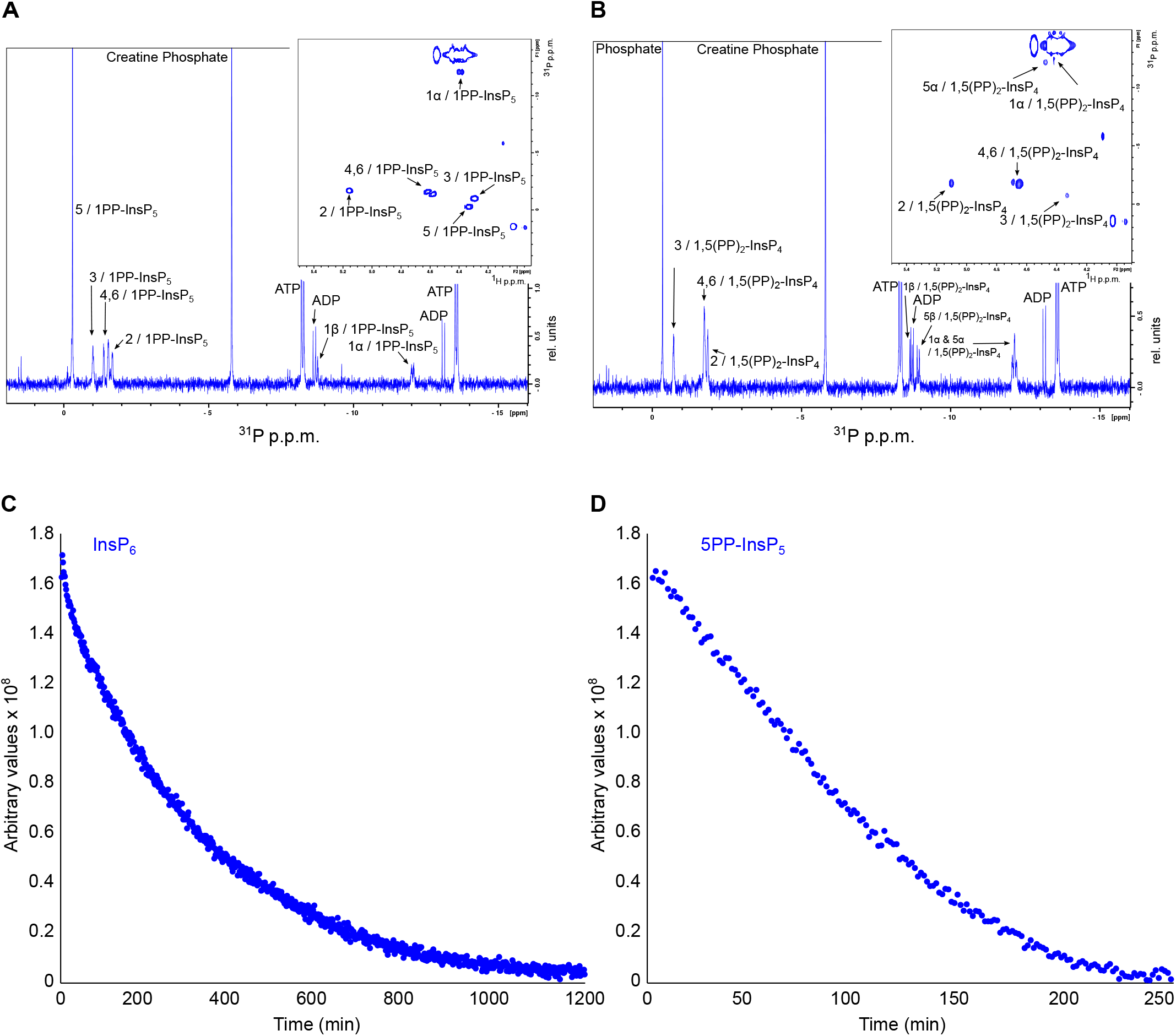
1D ^31^P and 2D ^1^H-^31^P-HMBC spectra of the products produced by AtVIH2-KD. **(A,B)** 1D ^31^P spectra of the products produced by the plant AtVIH2-KD in the presence of InsP_6_ (A) and 5PP-InsP_5_ (B). Inset panels shows the 2D ^1^H-^31^P-HMBC spectra. **(C, D)** Decay of the InsP_6_ (C) or 5-PP-InsP_5_ (D) substrate during the entire NMR-time course experiment.

**Figure 7 – figure supplement 1:**
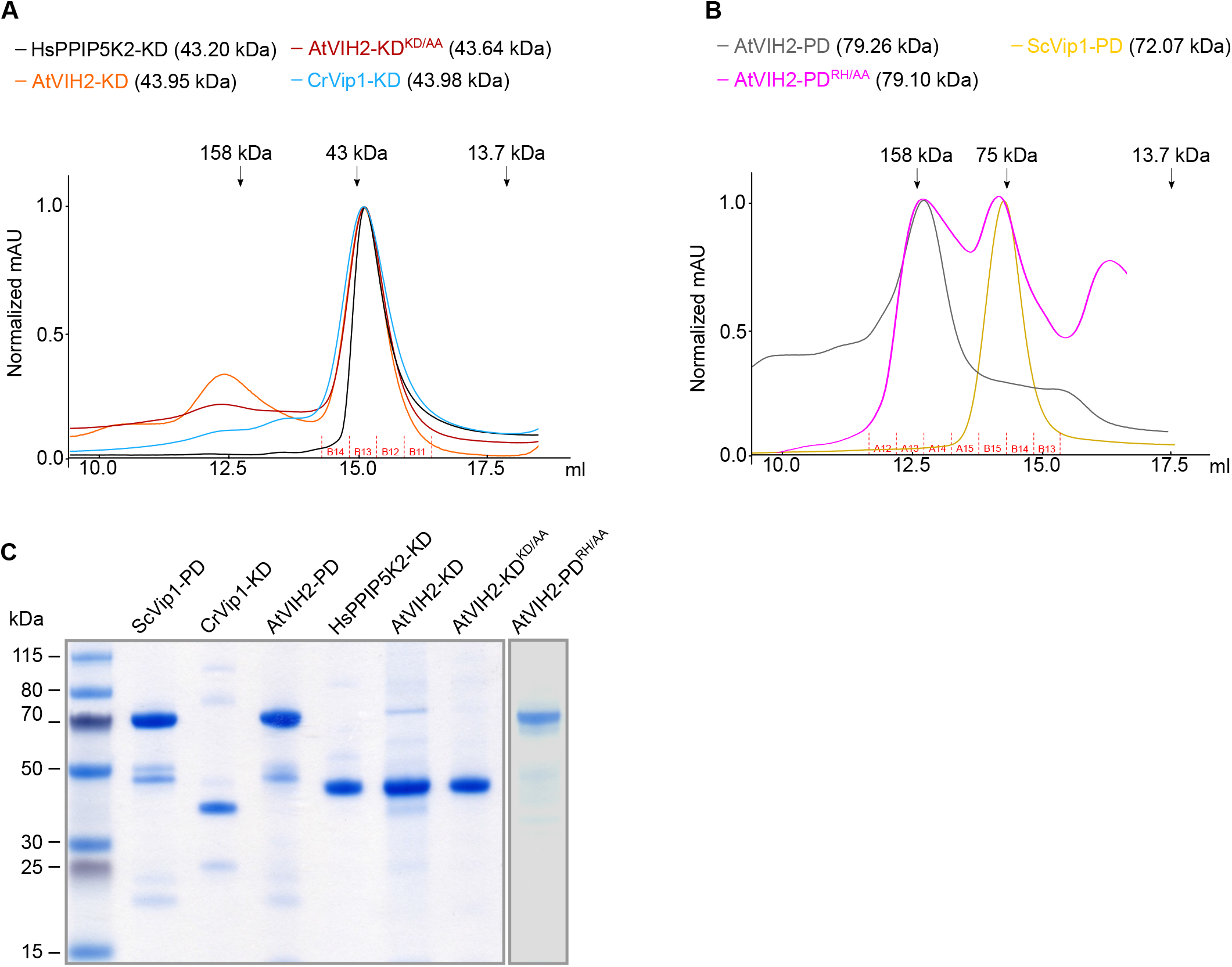
The purification of recombinant proteins. **(A)** Size-exclusion chromatography traces of purified recombinant HsVIP2 kinase domain (HsVIP2-KD, Glu41-Asp366), AtVIH2 kinase domain (AtVIH2-KD, Gly11-Asn338), AtVIH2 kinase domain mutant (AtVIH2-KD^KD/AA^, Gly11-Asn338), and CrVip1 kinase domain from Chlamydomonas (CrVip1-KD, Pro20-Ser350). Fractions isolated for enzymatic assays are indicated below. **(B)** Size-exclusion chromatography traces of purified recombinant AtVIH2 phosphatase domain (AtVIP2-PD, Gly359-Arg1002), AtVIH2 phosphatase domain mutant (AtVIP2-PD^RH/AA^, Gly359-Arg1002) and the ScVip1 phosphatase domain (ScVip1-PD, Ala515-Arg1088). For enzyme activity assays, the indicated fractions containing the purified recombinant protein were pooled (fractions A12-A15 for AtVIP2-PD; A12-B13 for AtVIP2-PD^RH/AA^; and A15-B13 for ScVip1-PD). **(C)** Coomassie-stained SDS-PAGE gel showing the recombinant proteins used in this study. For each lane, around 3 μg of protein from the respective concentrated fractions were loaded.

**Supplementary Table 1.** Transgenic lines used in this study.

| Promoter | Gene | Tag | Resistance | Genetic background |
| --- | --- | --- | --- | --- |
| *pVIH1* | *VIH1* | *mCitrine* | Basta | *vih1-2* |
| *pVIH2* | *VIH2* | *mCitrine* | Basta | *vih2-4* |
| *pVIH1* | *VIH1* | *mCitrine* | Basta | *vih1-2 vih2-4* |
| *pVIH2* | *VIH2* | *mCitrine* | Basta | *vih1-2 vih2-4* |
| *pVIH2* | *VIH2<sup>KD/AA</sup>* | *mCitrine* | Hygromycin | *vih1-2 vih2-4* |
| *pVIH2* | *VIH2<sup>RH/AA</sup>* | *mCitrine* | Hygromycin | *vih1-2 vih2-4* |
| *pVIH1* | *GUS* |  | Basta | Col Wt |
| *pVIH2* | *GUS* |  | Basta | Col Wt |
| *p35S* | *VIH1<sup>amiRNA</sup>* |  | Hygromycin | *vih2-4* |
| *pVIH2* | *H2B* | *mCitrine* | Basta | Col Wt |
| *pVIH2* | *H2B* | *mCitrine* | Basta | *vih2-4* |
| *pSPX1* | *H2B* | *eGFP* | Hygromycin | *spx1-1* |
| *pSPX3* | *H2B* | *eGFP* | Hygromycin | Col Wt |
| *pSPX1* | *SPX1* | *eGFP* | Hygromycin | *spx1-1* |
| *pSPX3* | *SPX3* | *eGFP* | Hygromycin | Col Wt |
| *pUbi10* | *VIH1* | *mCitrine* | Hygromycin | Col Wt |
| *pUbi10* | *VIH1<sup>KD/AA</sup>* | *mCitrine* | Hygromycin | Col Wt |
| *pUbi10* | *VIH1<sup>RH/AA</sup>* | *mCitrine* | Hygromycin | Col Wt |
| *pUbi10* | *VIH2* | *mCitrine* | Hygromycin | Col Wt |
| *pUbi10* | *VIH2<sup>KD/AA</sup>* | *mCitrine* | Hygromycin | Col Wt |
| *pUbi10* | *VIH2<sup>RH/AA</sup>* | *mCitrine* | Hygromycin | Col Wt |
| *pVIH2* | *HsVIP2* | *mCitrine* | Hygromycin | *VIH1(+/-) VIH2(-/-)* |
| *pVIH2* | *HsVIP2<sup>KD/AA</sup>* | *mCitrine* | Hygromycin | *VIH1(+/-) VIH2(-/-)* |
| *pVIH2* | *HsVIP2<sup>RH/AA</sup>* | *mCitrine* | Hygromycin | *vih1-2 vih2-4* |

**Supplementary Table 2.**
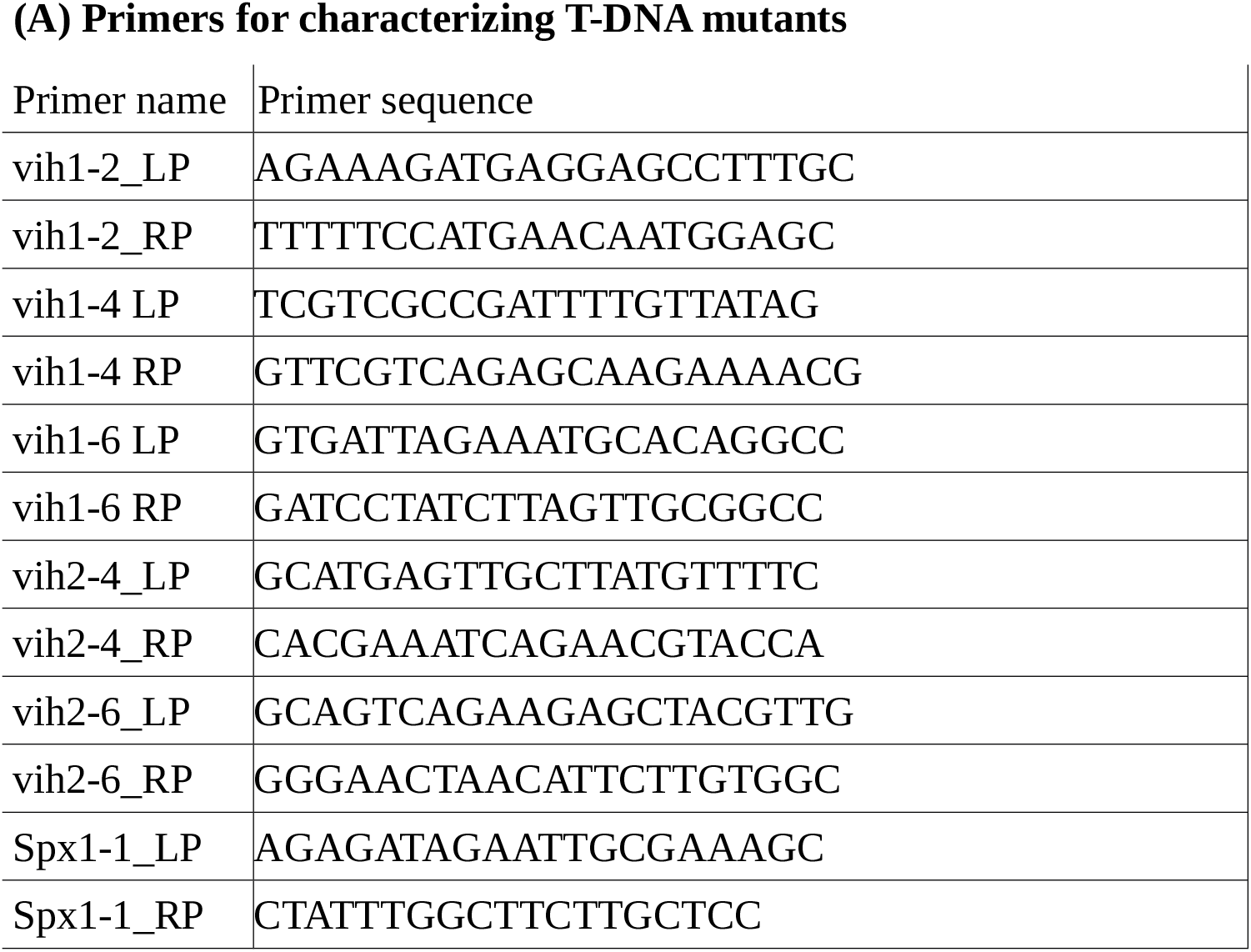
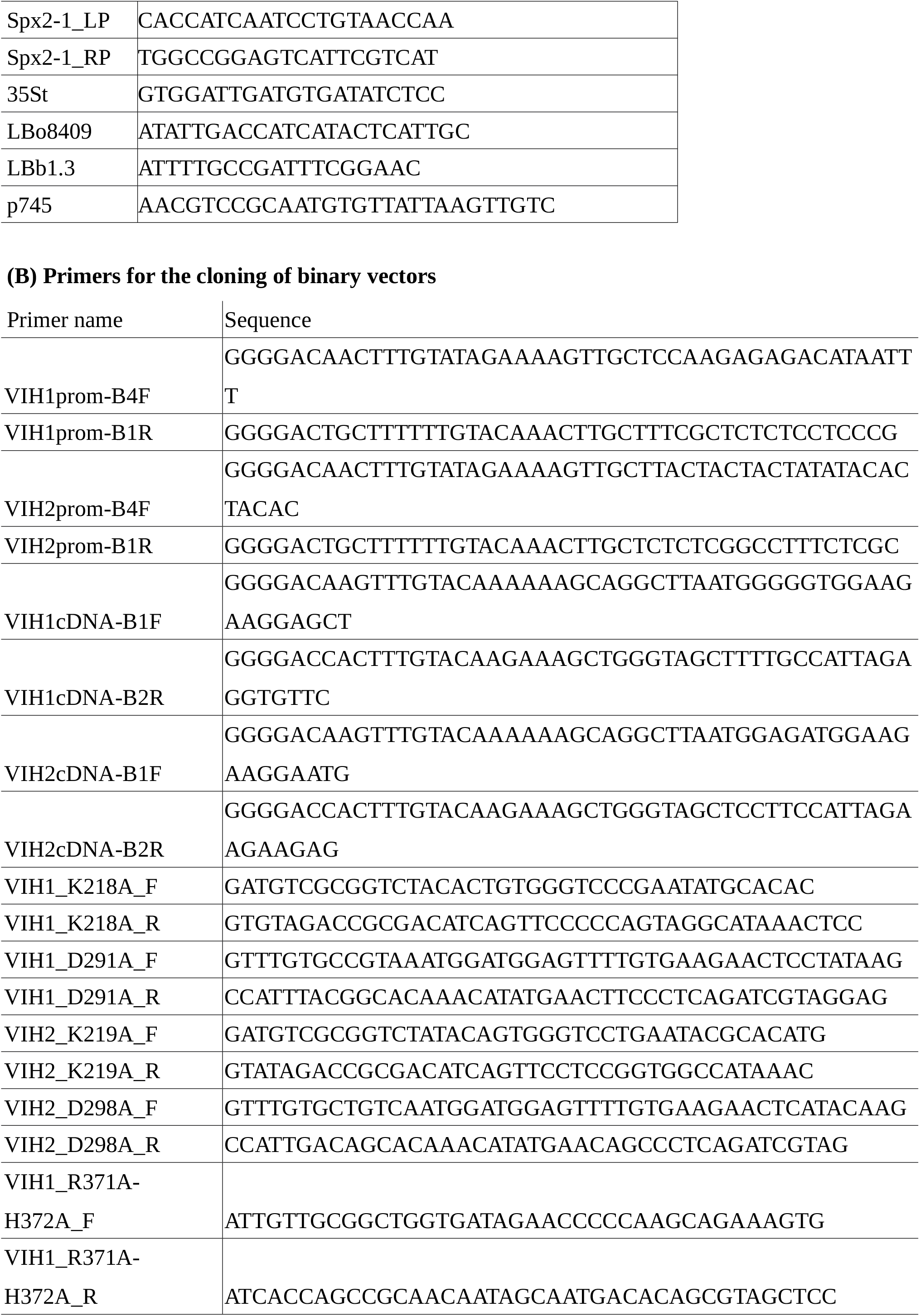
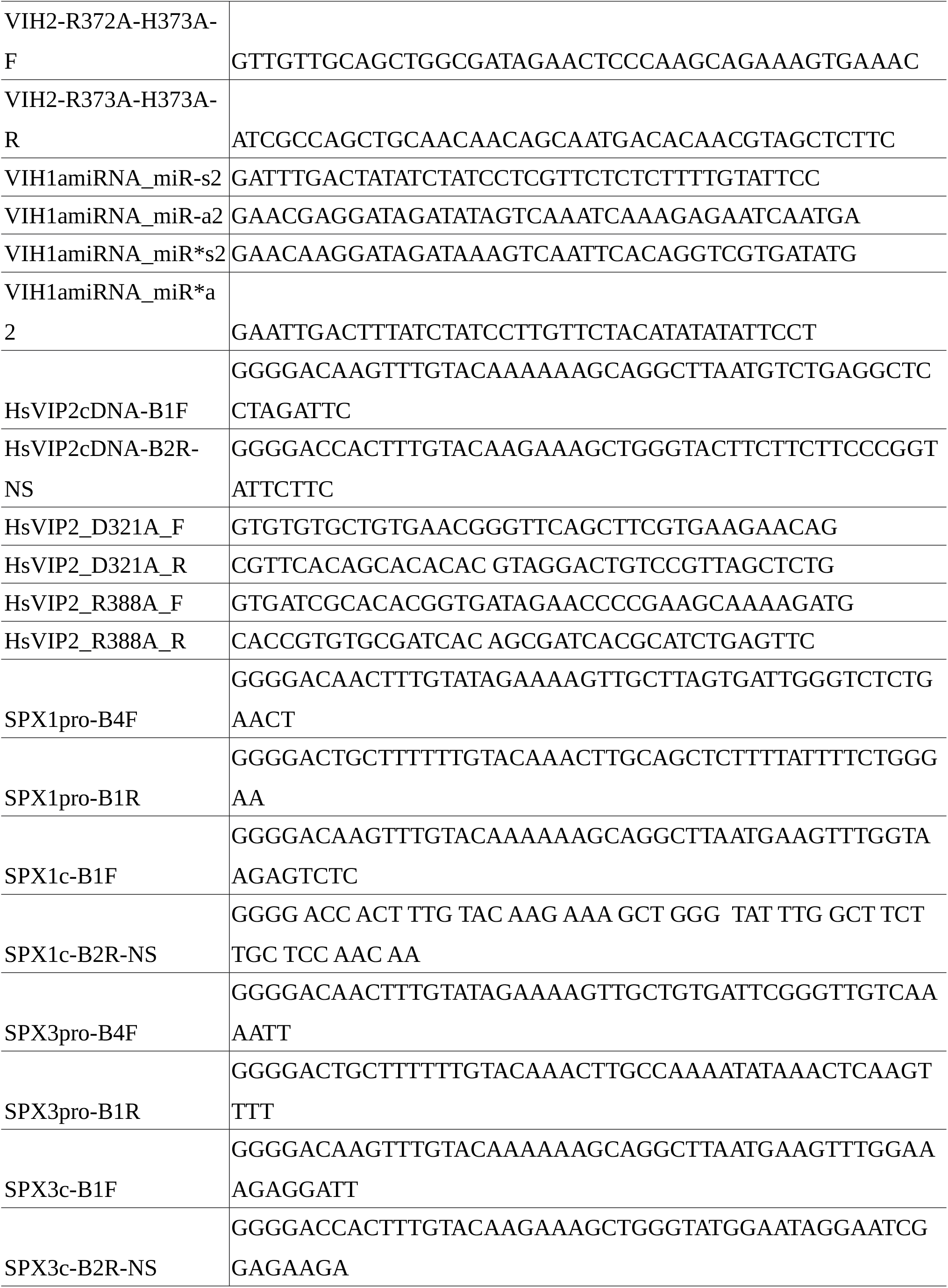
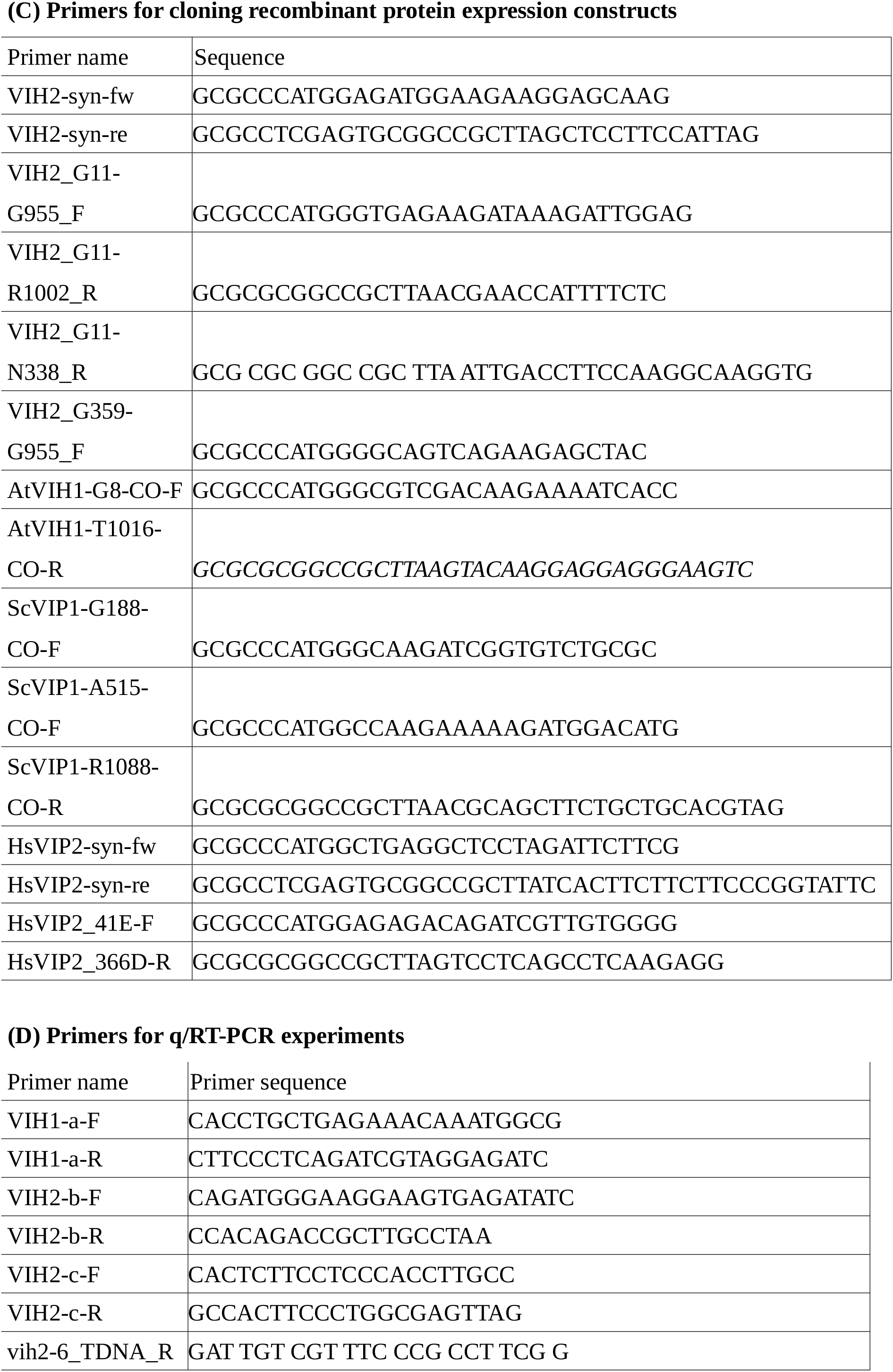
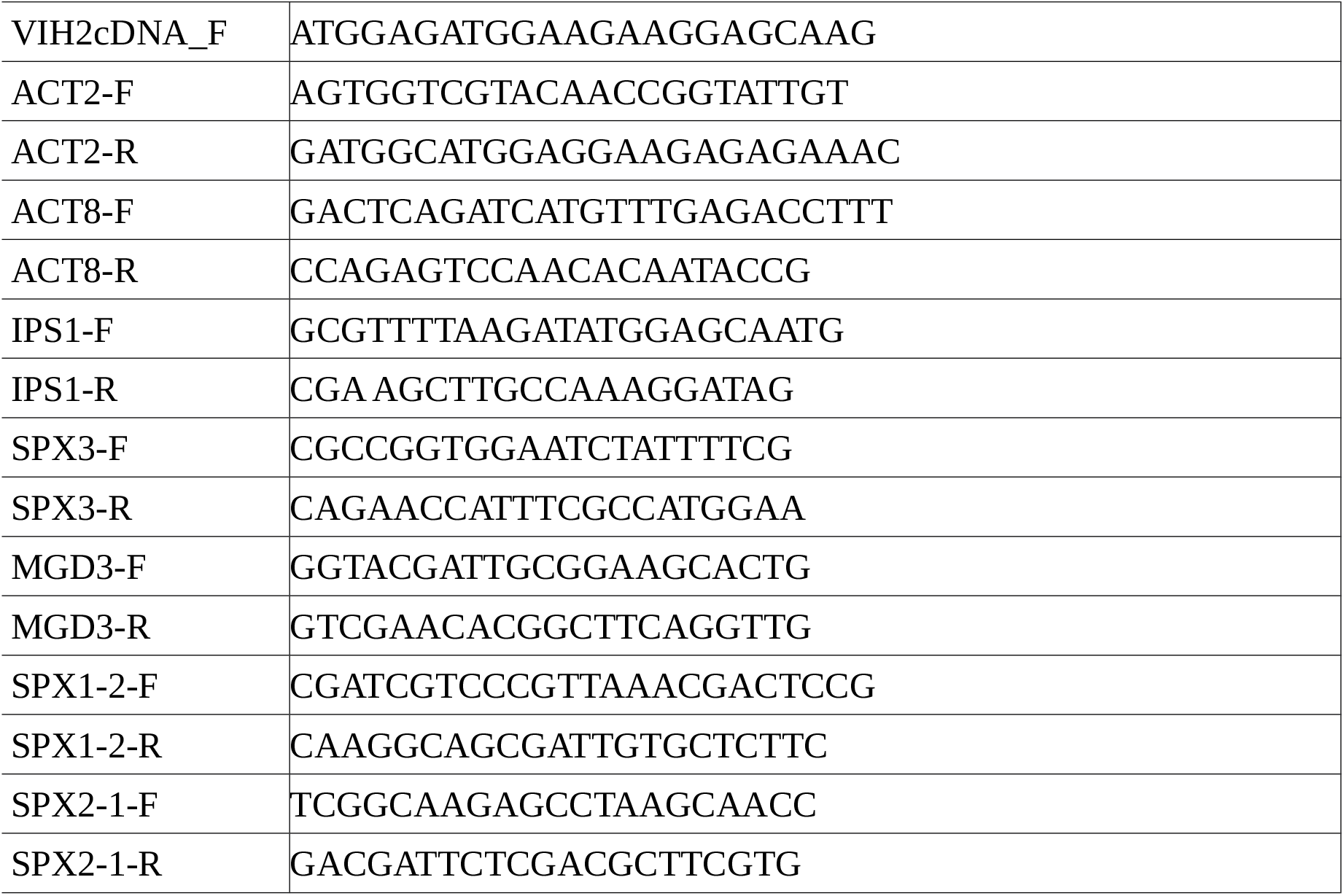
Primers used in this study.

**Supplementary Table 3.** The crossing between T-DNA lines and analysis of segregation ratio.

**(A) Over view of VIH double mutants**
|  | *vih1-2* ♀ | *vih1-4* ♀ | *vih1-6* ♀ | *vih1-2* ♂ | *vih1-4* ♂ | *vih1-6* ♂ |
| --- | --- | --- | --- | --- | --- | --- |
| *vih2-4* | D | W | D | D | W | D |
| *vih2-6* | W | W | W | W | W | W |

**(B) Analysis of segregation ratios in the progeny of independent lines of heterozygous plants**
|  | VIH1 (+/-); VIH2 (-/-) |  | VIH1 (-/-); VIH2 (+/-) |  |
| --- | --- | --- | --- | --- |
|  | D plants, % | W plants, % | D plants, % | W plants, % |
| #1 | 18.6 | 81.4 | 22.2 | 77.8 |
| #2 | 22.7 | 77.3 | 21.3 | 78.7 |
| #3 | 21.7 | 78.3 | 17.1 | 82.9 |
| #4 | 15.0 | 85.0 | 20.8 | 79.2 |
| #5 | 29.1 | 70.9 | 17.7 | 72.3 |
| #6 | 17.3 | 82.7 | 22.4 | 77.6 |

